# Two C-terminal sequence variations determine differential neurotoxicity between human and mouse α-synuclein

**DOI:** 10.1101/700377

**Authors:** Natalie Landeck, Katherine E. Strathearn, Daniel Ysselstein, Kerstin Buck, Sayan Dutta, Siddhartha Banerjee, Zhengjian Lv, John D. Hulleman, Jagadish Hindupur, Li-Kai Lin, Sonal Padalkar, George P. McCabe, Lia A. Stanciu, Yuri L. Lyubchenko, Deniz Kirik, Jean-Christophess Rochet

## Abstract

α-Synuclein (aSyn) aggregation is thought to play a central role in neurodegenerative disorders termed synucleinopathies, including Parkinson’s disease (PD). Mouse aSyn contains a threonine residue at position 53 that mimics the human familial PD substitution A53T, yet in contrast to A53T patients, mice show no evidence of aSyn neuropathology even after aging. Here we studied the neurotoxicity of human A53T, mouse aSyn, and various human-mouse chimeras in cellular and *in vivo* models as well as their biochemical properties relevant to aSyn pathobiology. We report that mouse aSyn is less neurotoxic than the human A53T variant as a result of inhibitory effects of two C-terminal amino acid substitutions on membrane-induced aSyn aggregation and aSyn-mediated vesicle permeabilization. Our findings highlight the importance of membrane-induced self-assembly in aSyn neurotoxicity and suggest that inhibiting this process by targeting the C-terminal domain could slow neurodegeneration in PD and other synucleinopathy disorders.

## Introductions

Parkinson’s disease (PD) is a common progressive neurodegenerative disorder characterized clinically by motor symptoms attributed to a loss of dopaminergic neurons in the *substantia nigra* (SN). At post-mortem examination neurons in various brain regions of PD patients present with cytosolic inclusions named Lewy bodies that contain amyloid-like fibrils of the presynaptic protein α-synuclein (aSyn) (1). A number of patients with familial forms of PD have been found to harbor mutations in the SNCA gene, including point mutations encoding the substitutions A30P, E46K, H50Q, G51D, A53E, and A53T and gene multiplications (2). Genetic and neuropathological findings in humans and data from animal model studies suggest that aSyn self-assembly plays a central role in the pathogenesis of PD and other neurodegenerative disorders involving an accumulation of aSyn aggregates in the brain, collectively referred to as synucleinopathies (3). A detailed understanding of molecular mechanisms by which aSyn forms neurotoxic aggregates is critical for developing therapies aimed at slowing neurodegeneration in the brains of patients with PD and other synucleinopathy disorders.

aSyn has been reported to adopt a natively unfolded, monomeric structure in solution (4) and to exist as a compact disordered monomer in mammalian cells (5), although additional evidence suggests that it can also exist as an oligomer in the cytosol (6). The protein is typically expressed as a 14.4 kDa polypeptide and consists of 3 domains. The N-terminal domain spanning residues 1-67 contains five conserved, lysine-rich repeats. The central region spanning residues 61-95 contains a sixth lysine-rich repeat and is highly hydrophobic. A key feature of this region is the presence of a segment spanning residues 71-82 that is required for aSyn aggregation (7). The C-terminal region spanning residues 96-140 is enriched with proline and acidic residues and is thought to regulate aSyn aggregation through auto-inhibitory long-range interactions (8, 9), with electrostatic interactions mediated by the acidic residues playing a major role in increasing the fibrillization lag time (10). aSyn binds to anionic phospholipid vesicles by forming an amphipathic α-helix with varying lengths, including a short N-terminal helix spanning residues ∼1-25 and a longer helix spanning residues ∼1-97 encompassing both the N-terminal and central hydrophobic domains (11, 12). Although membrane binding apparently plays an important role in the normal function of aSyn related to regulation of synaptic vesicle trafficking (12, 13), the protein has also been shown to undergo accelerated aggregation when incubated in the presence of phospholipid vesicles at high protein:lipid ratios (14–16). aSyn aggregation at the membrane surface is likely stimulated by the exposure of hydrophobic residues as the membrane-bound protein shifts from the long-helix form to the short-helix form (11, 16), and by the fact that molecular interactions needed for aSyn self-assembly likely occur with a higher probability on the two dimensional surface of the lipid bilayer than in solution (17, 18). Evidence from our laboratory suggests that membrane-induced aSyn aggregation plays a key role in neurotoxicity (16), potentially via a mechanism involving membrane permeabilization (19–22).

An interesting observation regarding aSyn biology is the fact that mouse aSyn (m-aSyn) naturally contains a threonine residue at position 53 and is thus identical to the human familial PD mutant A53T (h-aSyn A53T) at this site (23). This observation raises the question of why h-aSyn A53T triggers accelerated neurodegeneration in the brains of familial PD patients, whereas even aged mice show no evidence of neurodegeneration or the presence of Lewy bodies (24). This paradox is particularly striking given that recombinant m-aSyn forms amyloid-like fibrils more rapidly than h-aSyn WT or h-aSyn A53T (25, 26), apparently as a result of the disruption of long-range interactions involving the C-terminal region and the N-terminal and central hydrophobic domains (27). In fact, however, m-aSyn differs from h-aSyn-A53T by six mismatches, one in the central hydrophobic domain (S87N) and five in the C-terminal region (L100M, N103G, A107Y, D121G and D122S) (Figure 1A), and each of these could contribute to the apparent difference in neurotoxicity between the two variants.

**Figure 1.**
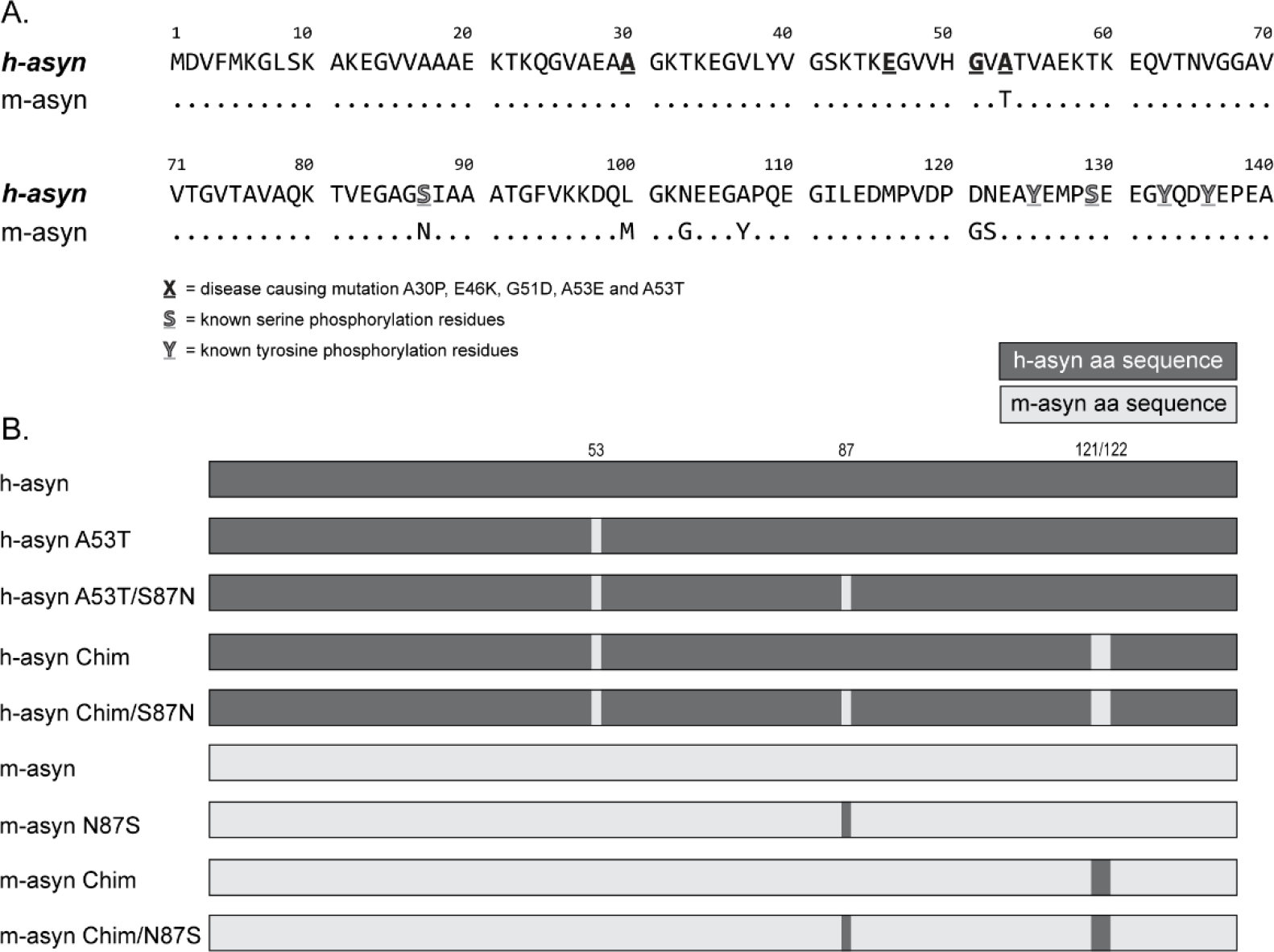
Human and mouse aSyn have seven sequence mismatches. (A) Sequence alignment of human, mouse, and chimeric aSyn variants showing mismatches between h-aSyn WT and m-aSyn at positions 53, 87, 100, 103, 107, 121, and 122. Also highlighted are the positions of familial PD substitutions and known serine and tyrosine phosphorylation sites. (B) Schematic representation of human, mouse, and chimeric aSyn variants examined in this study.

The goal of this study was to investigate the impact of the mismatches between m-aSyn and h-aSyn A53T on aSyn neurotoxicity. Previously it was reported that rat aSyn (which differs from the mouse protein by a single G-to-S mismatch at position 121) failed to trigger neuronal cell death when expressed from a lentiviral vector infused in rat SN, in contrast to h-aSyn WT and h-aSyn A53T (28). Similarly, we found that rat aSyn had a reduced ability to trigger nigral dopaminergic cell death compared to h-aSyn A53T in the rAAV-mediated overexpression model in the rat (29). Based on the close sequence similarity between mouse and rat aSyn, we hypothesized that the mouse protein is less neurotoxic than h-aSyn A53T, potentially because the toxic consequences of having a threonine residue at position 53 are alleviated by mismatch residues elsewhere in the murine sequence. To address this hypothesis, we compared h-aSyn A53T, m-aSyn, and various human-mouse chimeras in terms of their ability to elicit neurodegeneration in cellular and animal models relevant to PD and other synucleinopathy disorders. The human and mouse proteins and a subset of chimeras were further examined for their ability to form amyloid-like fibrils in the absence of membranes, undergo membrane-induced aSyn aggregation, and trigger membrane permeabilization.

## Results

### Neurotoxicity of aSyn variants in primary midbrain culture

In the first part of our study, we used a primary mesencephalic cell culture model to examine the effects of amino acid variations between m-aSyn and h-aSyn A53T on aSyn neurotoxicity. In addition to h-aSyn A53T and m-aSyn, two chimeric variants were analyzed: (i) h-aSyn Chimera, consisting of h-aSyn with the human-to-mouse substitutions A53T, D121G, and N122S; and (ii) m-aSyn Chimera, consisting of m-aSyn with the mouse-to-human substitutions G121D and S122N (Figure 1B). We chose to examine the effects of substitutions at positions 121 and 122 based on data showing that post-translational modifications near these residues, including C-terminal truncation and Y125 phosphorylation, modulate the protein’s aggregation propensity and neurotoxicity (30–34).

Primary midbrain cultures were processed as untreated control or transduced with adenovirus encoding each of the aSyn variants at multiplicities of infection (MOIs) that were adjusted to ensure equal expression of the different variants. We have previously shown that the adenoviral transduction efficiency is >90% for dopaminergic and non-dopaminergic neurons (16). The cells were fixed and stained for microtubule associated protein 2 (MAP2), a general neuronal marker, and tyrosine hydroxylase (TH), a marker of dopaminergic neurons. Relative dopaminergic cell viability was evaluated by determining the percentage of MAP2^+^ neurons that were also TH^+^. We found a significant reduction in the percentage of TH^+^ neurons in cultures transduced with h-aSyn A53T virus compared to untreated cultures or cultures expressing m-aSyn (Figure 2A, B). Cultures expressing m-aSyn Chimera exhibited a decrease in relative TH^+^ cell viability, similar to A53T-expressing cultures (Figure 2B). In contrast, h-aSyn Chimera had no impact on dopaminergic cell survival, similar to m-aSyn (Figure 2A). In additional experiments we found that cultures expressing A53T/D121G (but not A53T/N122S) exhibited a significant decrease in TH^+^ neuron loss compared to cultures expressing A53T (Figure 2C). Together these results suggested that (i) h-aSyn A53T is more neurotoxic than m-aSyn, and (ii) the human-to-mouse substitutions D121G and N122S (and in particular D121G) are at least partially responsible for the decrease in neurotoxicity of m-aSyn compared to h-aSyn A53T.

**Figure 2.**
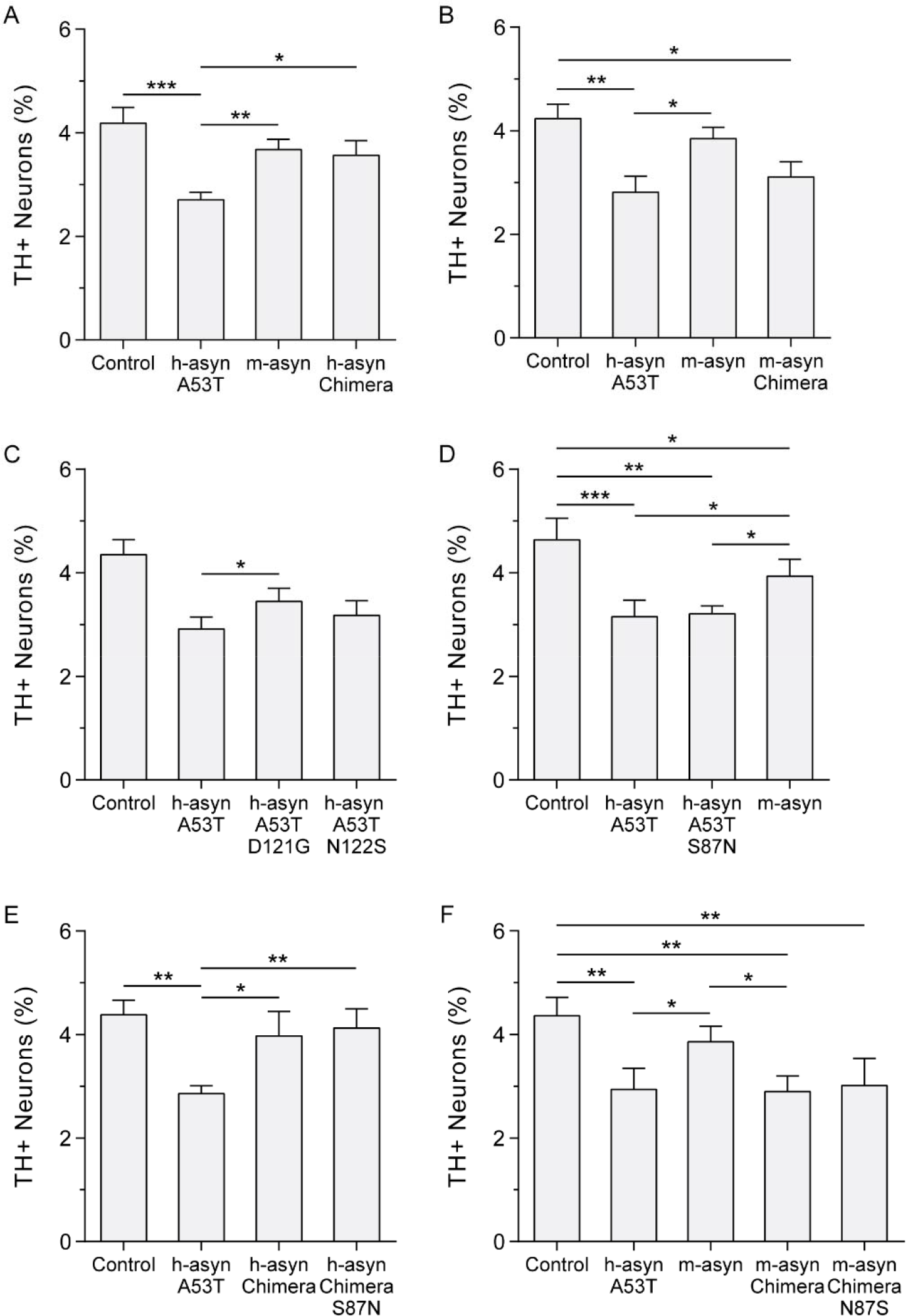
Human/mouse mismatches at positions 121 and 122 affect aSyn neurotoxicity in cell culture. Primary midbrain cultures were transduced with adenovirus encoding human, mouse, and chimeric aSyn variants at MOIs adjusted to ensure equal expression levels. Additional cultures were untransduced (‘control’). The cells were fixed, stained with antibodies specific for MAP2 and TH, and scored for dopaminergic cell viability. The data are presented as the mean ± SEM, n = 4 (A, B), n = 5 (C), or n = 3 (D-F). *p<0.05, **p<0.01, ***p<0.001; square root transformation, one-way ANOVA followed by Tukey’s multiple comparisons *post hoc* test.

In an independent set of experiments, we examined the impact of the mismatch between m-aSyn and h-aSyn A53T at position 87 (Figure 1) on aSyn-mediated dopaminergic cell death. Here our rationale was that residue 87 is located in the central hydrophobic domain, and substitutions and deletions in this region have been shown to modulate aSyn aggregation and neurotoxicity (7, 31). Cultures expressing h-aSyn A53T/S87N, h-aSyn Chimera S87N, or m-aSyn Chimera N87S exhibited no difference in relative dopaminergic cell viability compared to cultures expressing h-aSyn A53T (Figure 2D), h-aSyn Chimera (Figure 2E), or m-aSyn Chimera (Figure 2F), respectively. From these results, we inferred that the mismatch between m-aSyn and h-aSyn A53T at position 87 does not affect aSyn neurotoxicity.

Taken together, these findings suggest that residues D121 and N122 are important for the ability of h-aSyn A53T to elicit dopaminergic cell death. Moreover, we infer that residues G121 and S122 (but not N87) of m-aSyn interfere with the expected neurotoxic effect of the threonine residue at position 53 of the mouse sequence.

### Neurotoxicity of aSyn variants in rat nigrostriatal projection neurons

To further examine the effects of mismatches at positions 121 and 122 of m-aSyn and h-aSyn A53T on aSyn neurotoxicity, we characterized h-aSyn A53T, m-aSyn, h-aSyn Chimera, and m-aSyn Chimera in terms of their toxicity to nigral dopamine neurons *in vivo*. rAAV vectors encoding these variants were injected unilaterally into the rat SN at two different titers, corresponding to a low-toxicity titer at 1.3E13 gc/mL (n=4-6 per group) and a high-toxicity titer at 3.5E14 gc/mL (n=4-10 per group). Animals were killed 8 weeks after transduction, and serial coronal midbrain sections were stained for histological analysis to assess transgene expression and the extent of neurodegeneration in the nigral dopamine neurons. Representative specimens of animals injected with a high-titer vector are shown in Figure 3A-Y. The aSyn antibody applied here was used to evaluate transgene expression and is specific for aSyn but does not distinguish between human and rodent variants as seen in Figure 3A, where endogenous aSyn can be visualized especially in the axon terminal located in the SN pars reticulata contralateral to the AAV injection. On the injected side, transduced cells over-expressing h-aSyn A53T (Figure 3B), h-aSyn Chimera (Figure 3C), m-aSyn (Figure 3D), or m-aSyn Chimera (Figure 3E) were visible in the SN pars compacta. The human aSyn-specific syn211 antibody resulted in specific staining in groups expressing h-aSyn A53T or m-aSyn Chimera, whereas the contralateral side of rat midbrain sections as well as in sections expressing m-aSyn or h-aSyn Chimera were devoid of any immunoreactivity (Figure S1). Notably, staining with a pan aSyn antibody, Syn-1, showed that the overexpression was present in all groups, suggesting that the affinity of syn211 to the aSyn variants was dependent on the identity of the residues at positions 121 and 122 (Figure S1F-J).

**Figure 3.**
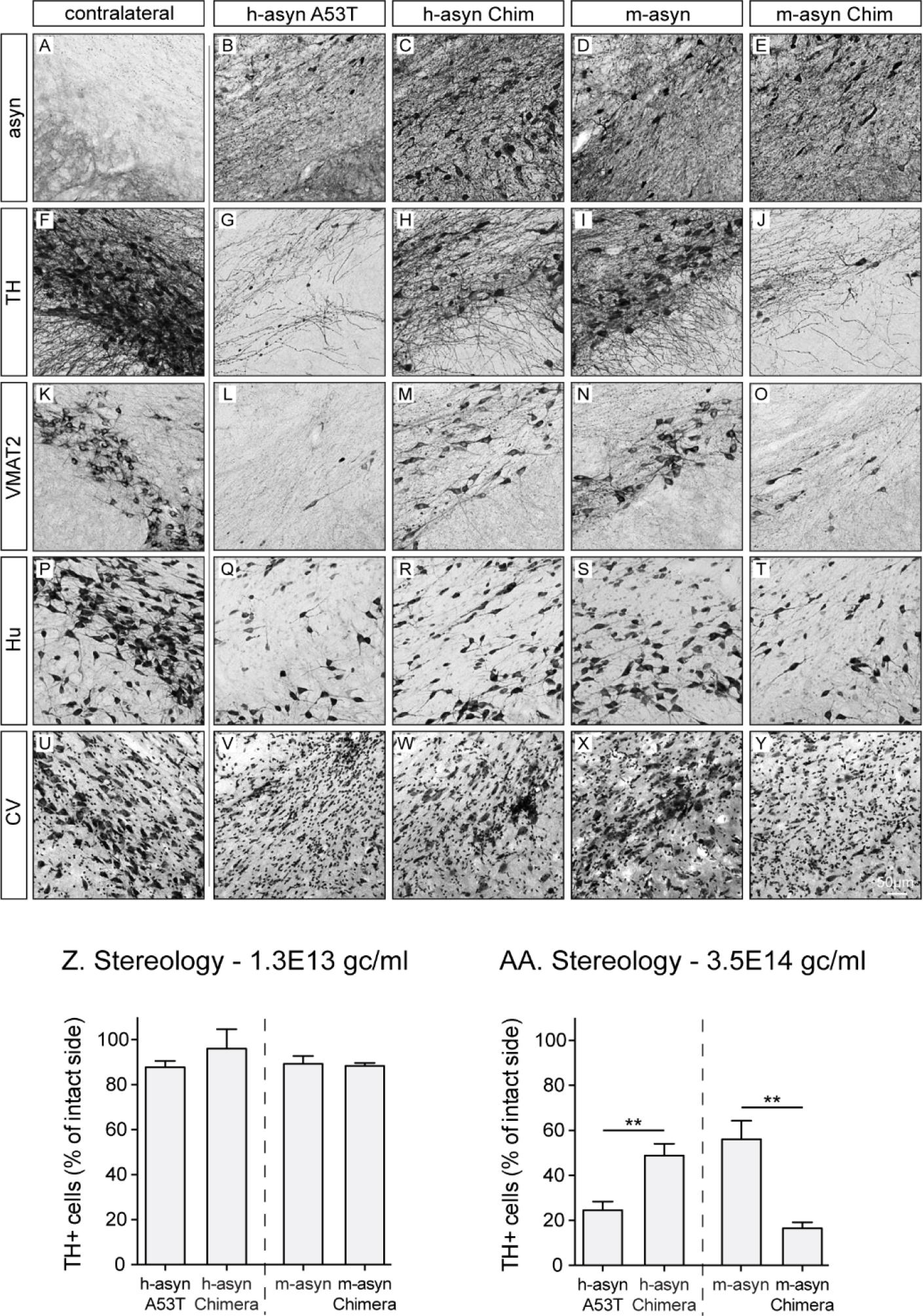
Human/mouse mismatches at positions 121 and 122 affect aSyn-mediated dopaminergic cell death in rat midbrain. Rats (4-10 per group) were injected unilaterally into the SN with vectors encoding h-aSyn A53T, h-aSyn Chimera, m-aSyn, or m-aSyn Chimera at two different titers (1.3E13 gc/ml or 3.5E14 gc/ml). Eight weeks after surgery, coronal midbrain sections of the high-dose group were stained for aSyn in order to visualize transgene expression (A-E). Adjacent specimens were stained for TH (F-J), VMAT2 (K-O), Hu (P-T) or cresyl violet (U-Y). The first panel in each row shows a representative image of the contralateral uninjected side (A, F, K, P, U). Counts of TH+ nigral dopaminergic neurons are shown in Z for the lower vector dose and in AA for the higher vector dose. Data are presented as the percentage of intact side cell numbers ± SEM. **p<0.01; two-tailed Mann-Whitney U test. Scale bar, 50 μm (panels A-Y). *See also Figure S1*.

Immunohistochemical analyses were carried out to evaluate the loss of dopaminergic neurons in rat midbrain sections stained for TH (Figure 3F-J) and vesicular monoamine transporter 2 (VMAT2) (Figure 3K-O). A clear loss of TH^+^ as well as VMAT2^+^ neurons was seen in all groups when compared to the uninjected contralateral side (Figures 3F and 3K, respectively). The reduction in cell numbers was not due to a loss of expression of dopaminergic markers, as it was clearly visible also in specimens stained with antibodies against the neuronal marker proteins Hu (Figure 3P-T) or with cresyl violet (CV) (Figure 3U-Y). In order to quantify dopaminergic cell loss in the SN, we counted TH^+^ cells in serial sections from both the low-titer and high-titer vector groups. Results obtained for the low-titer group showed no cell death for any of the aSyn variants and no significant differences among groups (Figure 3Z). In contrast, in the 3.5E14 gc/mL titer groups, the expression of h-aSyn A53T resulted in a significantly higher percentage of cell death when compared to the h-aSyn Chimera group (75 ± 4 versus 50 ± 5% cell loss, respectively; Figure 3AA). Moreover, the m-aSyn group displayed significantly reduced neurotoxicity in TH^+^ neurons than the m-aSyn Chimera group (56 ± 8 versus 17 ± 3% cell survival, respectively). These results suggested a considerable effect of the human/mouse mismatches at positions 121 and 122 on aSyn-mediated dopaminergic cell death in rat SN.

To determine whether the significant differences in dopaminergic cell death observed in the SN were also evident in the nigrostriatal projections, we evaluated the TH^+^ and VMAT2^+^ fiber intensity in the striatum of the high titer groups. In serial sections throughout the rostocaudal axis of the striatum, we found a reduction in staining intensity for both TH (Figure 4A-E, A’-E’) and VMAT2 (Figure 4G-K, G’-K’) for all groups when compared to contralateral striatal areas. A significant reduction of innervation density was observed for the m-aSyn Chimera group when compared to the m-aSyn group for both TH^+^ (Figure 4F) and VMAT2^+^ projections (Figure 4L). These observations are in accordance with dopaminergic cell loss seen in the corresponding SN sections of the same animals. There was a trend for a difference in the h-aSyn Chimera group when compared to the h-aSyn A53T group, although this did not reach significance [TH (p=0.11); VMAT2 (p=0.27)].

**Figure 4.**
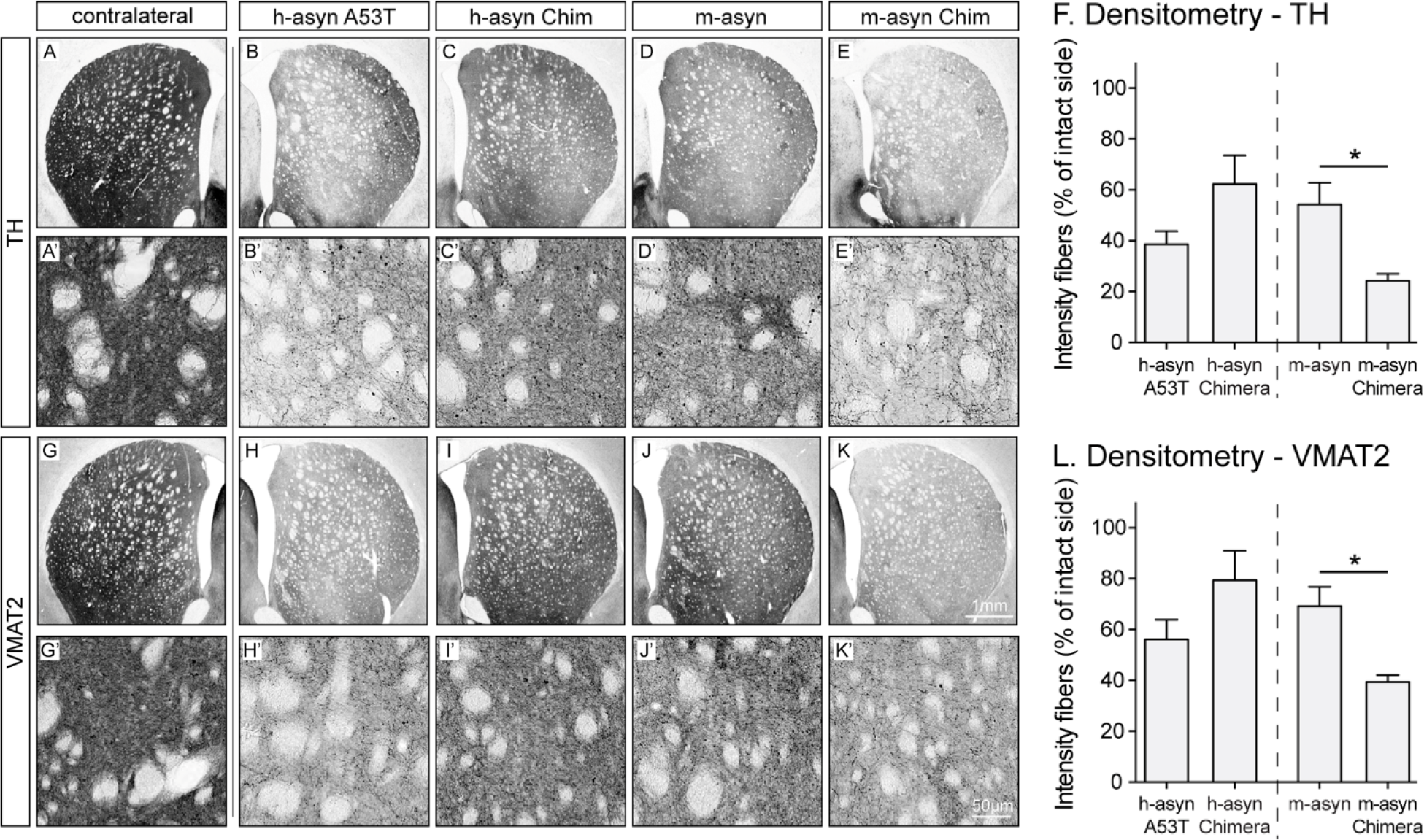
Mouse-to-human substitutions at positions 121 and 122 enhance the ability of m-aSyn to elicit dopaminergic fiber loss in rat striatum. Representative striatal rat brain sections stained for TH or VMAT2 (4-6 per group) are shown in panels A-E or G-K, respectively, as low-magnification images, and in panels A’-E’ or G’-K’, respectively, as corresponding high-magnification images. The imaged sections were from animals injected with the highest vector dose. The first panel of each row shows a representative image of the contralateral uninjected side (A, A’, G, G’). Quantified staining intensities of the striatum are displayed in F for TH and L for VMAT2. Data are presented as the percentage of corresponding contralateral staining intensity ± SEM. *p<0.05; two-tailed Mann-Whitney U test. Scale bar, 1 mm (panels A-K) or 50 μm (panels A’-K’). *See also Figure S2*.

### Aggregate formation by aSyn variants in rat midbrain

To determine whether the aSyn variants differed in terms of aggregation propensity in rat midbrain, we treated striatal sections with Proteinase K (PK) to digest non-aggregated species and reveal digestion-resistant aSyn assemblies. As a reference for the PK incubation we chose the endogenous aSyn staining in the CA3 region of the hippocampus (Figure S2A, F, K), which would not be affected by nigral transduction. Whereas untreated striatal sections stained for aSyn showed both punctate and fiber-like staining (Figure S2B-E), a clearance of digested and therefore non-aggregated aSyn could already be appreciated after 5 min of PK incubation (Figure S2F-J). At a later incubation time (45 min) when endogenous aSyn staining could no longer be detected in the CA3 region (Figure S2K), and therefore all non-aggregated synuclein would be digested, resistant aSyn aggregates were detected in striatal sections in all groups (Figure S2L-O). These results suggested that the human/mouse mismatches at positions 121 and 122 do not have a major impact on the ability of aSyn to form PK-resistant aggregates in the striatal fiber terminals of dopaminergic neurons.

### Effects of human/mouse aSyn mismatches on aSyn fibrillization rates

The next set of experiments was aimed at determining the effects of human/mouse aSyn mismatches on rates of aSyn fibrillization. Based on evidence that m-aSyn forms amyloid-like fibrils more rapidly than h-aSyn WT or A53T (25, 26), we hypothesized that a more rapid conversion of natively unfolded aSyn to mature, amyloid-like fibrils could lead to a depletion of toxic oligomers (26, 35, 36). As a corollary, we reasoned that the human-to-mouse substitutions could stimulate aSyn fibrillization, which in turn could account for the reduced neurotoxicity of m-aSyn and h-aSyn Chimera compared to h-aSyn A53T. To address this hypothesis, we monitored the fibrillization of human, mouse, and chimeric aSyn variants using a thioflavin T fluorescence assay. Consistent with previous results, we found that m-aSyn and h-aSyn A53T formed fibrils with a markedly reduced lag time compared to h-aSyn WT, although the decrease in lag time was more pronounced for the mouse protein (Figure 5A). Analysis of the chimeric aSyn variants revealed that h-aSyn Chimera formed fibrils with a longer lag time compared to h-aSyn A53T, whereas the rate of fibrillization was only slightly decreased for m-aSyn Chimera compared to m-aSyn (Figure 5B).

**Figure 5.**
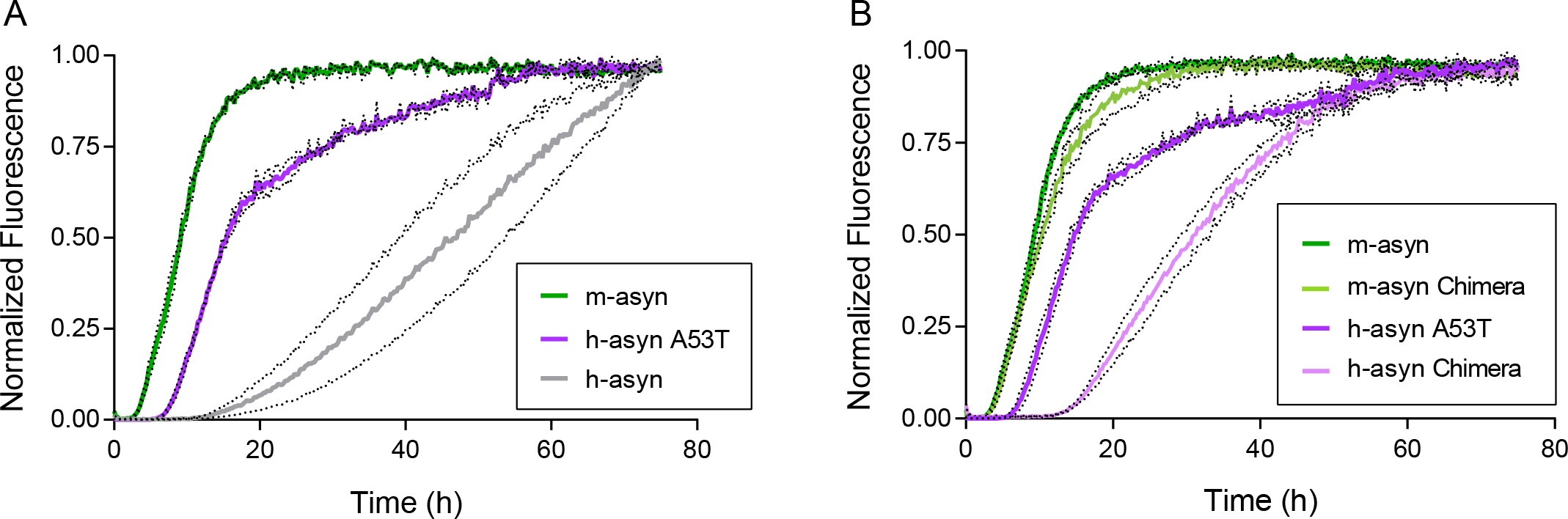
M-aSyn forms amyloid-like fibrils with a shorter lag time compared to human and chimeric aSyn variants. The formation of amyloid-like fibrils was monitored in solutions of monomeric h-aSyn WT, h-aSyn A53T, and m-aSyn (A), or h-aSyn A53T, h-aSyn Chimera, m-aSyn Chimera, and m-aSyn (B) (35 µM of each). The protein solutions were incubated at 37 °C with constant agitation and analyzed at various times for thioflavin T fluorescence. The graphs show the mean normalized fluorescence (determined from 3 independent experiments; 3 technical replicates per experiment) plotted against the incubation time. The dotted lines above and below each solid line correspond to the mean signal ± SEM. *See also Figure S3*.

Secondly, we investigated the effects of the human/mouse mismatch at position 87 on fibrillization rates, based on the premise that residue 87 is in the central hydrophobic region, and substitutions or deletions in this region have been shown to modulate aSyn self-assembly (7, 31). In contrast to our observation that substitutions at position 87 had no impact on aSyn neurotoxicity in primary midbrain cultures (Figure 2), A53T/S87N and h-aSyn Chimera/S87N exhibited a reduced fibrillization lag time compared to h-aSyn A53T and h-aSyn Chimera, respectively (Figure S3A, B). Moreover, m-aSyn N87S and m-aSyn Chimera/N87S formed amyloid-like fibrils with a longer lag time compared to m-aSyn and m-aSyn Chimera, respectively (Figure S3C, D).

Collectively, these results suggest that differences in neurotoxicity among the human, mouse, and chimeric aSyn variants (Figures 2-4) correlate weakly with differences in fibrillization rates.

### Effects of human/mouse aSyn mismatches on fibril morphology and dimensions

Next, we examined the impact of human/mouse aSyn mismatches on fibril morphology and dimensions based on evidence that different types of aSyn fibrils (e.g. cylindrical fibrils versus flatter ribbons) can trigger different levels of neurotoxicity in cellular and animal models (37). Examination of incubated samples of human, mouse, and chimeric aSyn by atomic force microscopy (AFM) revealed that all of the variants formed straight, rigid, unbranched fibrils, organized either as individual fibrils or in fibril bundles (Figure 6A-D). Two types of fibrils (‘twisted’ and ‘non-twisted’) were detected in samples of m-aSyn and h-aSyn-Chimera, whereas only non-twisted fibrils were observed in samples of h-aSyn-A53T and m-aSyn-Chimera under our experimental conditions. The heights of the non-twisted fibrils ranged from ∼8 to ∼11 nm, whereas the heights of the ‘valley’ and ‘peak’ regions of the twisted fibrils ranged from ∼7 to ∼9 nm and ∼11 to ∼12 nm, respectively. The average pitch of twisted fibrils formed by m-aSyn and h-aSyn Chimera was 123 ± 5 nm and 91 ± 8 nm, respectively (Figure 6E,F).

**Figure 6.**
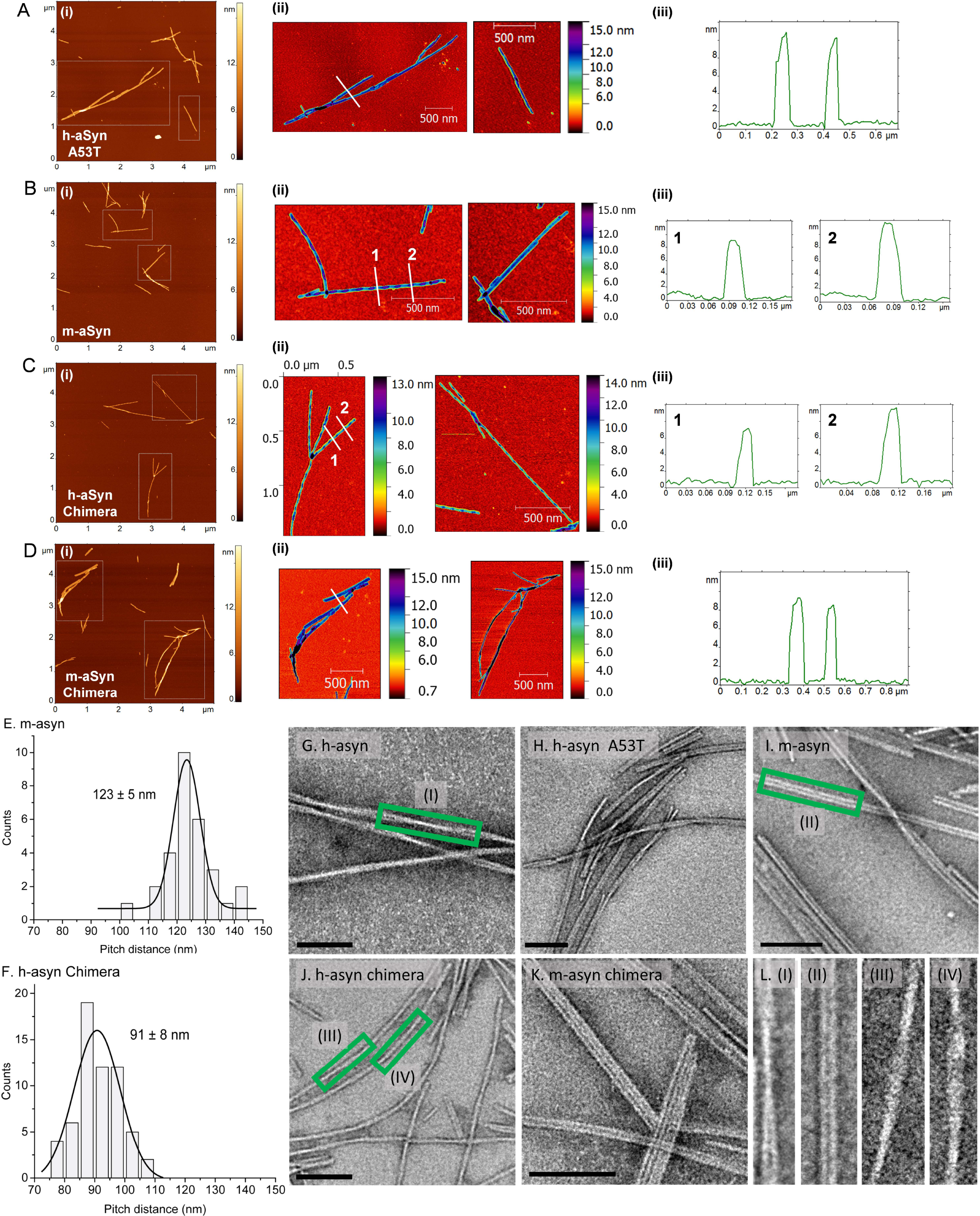
Human, mouse, and chimeric aSyn variants form amyloid-like fibrils with similar dimensions but different morphologies. (A-D) Results of AFM analysis of fibrils prepared from recombinant h-aSyn A53T (A), m-aSyn (B), h-aSyn Chimera (C), and m-aSyn Chimera (D). Panels A(i) – D(i) on the left show typical AFM topographic images of fibrils formed by each aSyn variant. Dashed boxes indicate areas shown at a higher magnification in a spectral color scheme in panels A(ii) – D(ii). Panels A(iii)-D(iii) on the right show cross-section profiles. Regions of the fibrils from which the profiles are taken are shown with white solid lines on the AFM images in panels A(ii)-D(ii). Twisted fibrils were observed for m-aSyn (B) and h-aSyn Chimera (C). Twisted and non-twisted fibrils account for 65% and 35% (respectively) of total fibrils formed by m-aSyn, and 76% and 24% (respectively) of total fibrils formed by h-aSyn Chimera. Cross-section profiles in panels A(iii) and D(iii) reveal heights of non-twisted fibrils (∼8-11 nm). Cross-section profiles 1 and 2 reveal heights of twisted fibrils at valley regions (∼7-9 nm) and peak regions (∼11-12 nm), respectively, in panels B(iii) and C(iii). (E, F) Histograms showing distances between peaks of twisted fibrils formed by m-aSyn (E) and h-aSyn Chimera (F). The distances were determined from cross-section profiles obtained along the long axis of the fibrils. (G-L) Representative TEM images of fibrils formed by h-aSyn WT (G), h-aSyn A53T (H), m-aSyn (I), h-aSyn Chimera (J), and m-aSyn Chimera (K). Scale bar, 100 nm. (L) Higher-magnification images generated from boxed regions of panels G, I, and J show a pair of fibrils wound around each other via a helical twist (I), two fibrils aligned in parallel (II), or individual fibrils with a twisted morphology (III and IV).

Further analysis of the fibrils by TEM also revealed the presence of straight, rigid, unbranched fibrils for all of the variants (Figure 6G-L). The fibrils had similar properties as those described for previously reported aSyn amyloid-like fibrils (e.g. diameter of ∼10-15 nm; apparent winding of two fibrils around each other via a helical twist) (26). Similar features were also observed for fibrils formed by aSyn variants with the S87N or N87S substitution. There were no striking differences in fibril morphology among the different aSyn variants, except that individual fibrils formed by h-aSyn-Chimera had a twisted appearance similar to that observed for this variant by AFM (in contrast, twisted fibrils of m-aSyn were not observed by TEM). It is unclear why different results were obtained for m-aSyn and h-aSyn Chimera via AFM versus TEM, although this discrepancy can potentially be explained by the fact that (i) the variants formed two different types of twisted fibrils, as implied by their different pitch values (Figure 6E,F), and (ii) different types of fibrils adsorb with different efficiencies to substrates with different properties such as the hydrophilic mica and hydrophobic carbon-coated grids used here for AFM and TEM, respectively (38).

Overall, these data suggest that differences in the morphologies of aSyn fibrils formed under the self-assembly conditions used in this study could potentially contribute to differences in neurotoxicity among the human, mouse, and chimeric aSyn variants (Figures 2-4).

### Effects of human/mouse aSyn mismatches on membrane-induced aSyn aggregation

Based on our earlier finding that lipid-induced self-assembly plays a key role in aSyn neurotoxicity (16), we hypothesized that human-to-mouse substitutions should lead to a decrease in aSyn aggregation propensity at membrane surfaces. Our rationale was that although aSyn aggregation is primarily driven by the central hydrophobic region (7, 11, 16), modifications to the C-terminal domain have been shown to modulate aSyn self-assembly (39) and aSyn-membrane interactions (40) through effects on long-range interactions.

As a first step, we characterized the human, mouse, and chimeric variants in terms of membrane affinities, based on the premise that differences in affinity could lead to differences in the variants’ propensity to undergo membrane-induced aggregation (16). Recombinant h-aSyn WT, h-aSyn A53T, h-aSyn Chimera, m-aSyn, and m-aSyn Chimera in a monomeric state were titrated with small unilamellar vesicles (SUVs) composed of egg phosphatidylglycerol (PG) and egg phosphatidylcholine (PC) (1:1 mol/mol). Analysis of the samples via far-UV CD, a method used to monitor the increase in aSyn α-helical structure that results from binding of the protein to phospholipids, yielded similar lipid titration curves, Kd values, and maximum α-helical contents for all of the aSyn variants (Figure 7A; Table S1). These results suggested that the C-terminal human/mouse mismatches have little effect on the affinity of aSyn for phospholipid membranes, or on the degree of α-helical structure adopted by the protein in the presence of saturating lipid.

**Figure 7.**
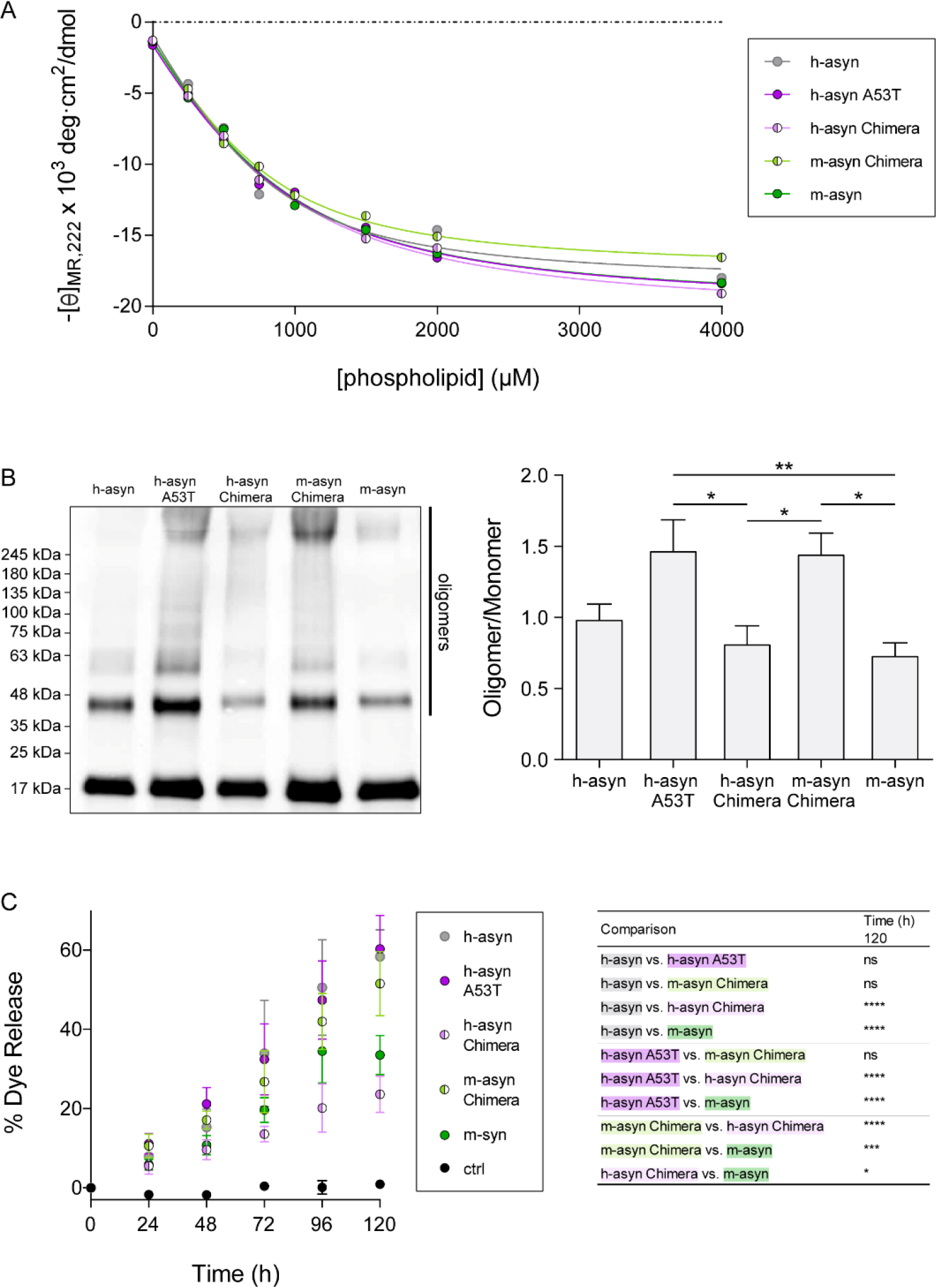
Human/mouse mismatches at positions 121 and 122 affect membrane-induced aSyn aggregation and aSyn-mediated vesicle disruption. (A) Solutions of recombinant aSyn variants were incubated with increasing concentrations of SUVs composed of egg PG:PC (1:1mol:mol) and analyzed by far-UV CD to determine [θ]_*MR*,222_. The data were fit to equation 3 (see‘Experimental Procedures’), and values for KD, minimum [θ]_*MR*,222_, and maximum α-helical content were determined from the values of the fit parameters (Table S1). (B) Monomeric aSyn variants were incubated with PG:PC SUVs (protein-to-lipid ratio, 1:20 mol/mol) at 37 °C for 72 h. Left: Representative Western blot image (the vertical line to the right of the image shows the region of the blot with bands corresponding to oligomeric forms of aSyn). Right: Bar graph showing the ratio of total oligomer band intensity to monomer band intensity determined for each sample. (C) Calcein-loaded SUVs were incubated in the absence (‘ctrl’) or presence of aSyn variants. Calcein release was monitored via fluorescence measurements using excitation and emission wavelengths of 485 nm and 515 nm, respectively. Left: graph showing % dye sssrelease versus time, with 100% release determined as the signal obtained from vesicles treated with Triton X-100. Right: table listing P values determined for vesicle permeabilization data obtained at the 120 h time point (a more complete listing of P values at different time points is presented in Table S2). The data in (B) and (C) are presented as the mean ± SEM, n = 4 (B) or n = 3 (C). *p<0.05, **p<0.01, ***p<0.001, ****p<0.0001, one-way ANOVA followed by Tukey’s multiple comparisons *post hoc* test (a square root transformation was carried out on the data in panel C).

Next, we compared the human, mouse, and chimeric aSyn variants in terms of their ability to undergo membrane-induced aggregation using a lipid-flotation assay. The proteins were incubated with PG:PC SUVs under conditions that promote aSyn self-assembly, and the membrane fraction was isolated by gradient centrifugation and analyzed via Western blotting. The lane loaded with h-aSyn WT contained immunoreactive bands at ∼45 and 60 kDa, whereas the lane containing h-aSyn A53T or m-aSyn Chimera displayed more extensive laddering, including bands at 75 and 100 kDa, and an intense smear of aSyn immunoreactivity in the region of the gel above 245 kDa (Figure 7B). In contrast, high molecular-weight species were markedly less abundant in the lanes loaded with h-aSyn Chimera or m-aSyn. Densitometry analysis revealed a ∼2-fold greater ratio of total oligomer band intensity to monomer band intensity in the h-aSyn A53T and m-aSyn Chimera samples compared to the h-aSyn Chimera and m-aSyn samples, respectively (Figure 7B). Although the results revealed a trend towards an increase in the oligomer/monomer ratios for h-aSyn A53T and m-aSyn Chimera compared to the human WT protein, these effects did not reach statistical significance. Collectively, these data suggested that the human-to-mouse substitutions at positions 121 and 122 interfere with membrane-induced aSyn aggregation.

### Effects of human/mouse aSyn mismatches on a-Syn-mediated membrane disruption

A number of groups have reported that aSyn aggregation at the membrane can affect bilayer integrity via a mechanism involving phospholipid extraction coupled with aggregate growth (19, 21). Recently, we found that two familial aSyn mutants with a high propensity to form oligomers at membrane surfaces (A30P and G51D) (16) efficiently triggered vesicle disruption under incubation conditions suitable for membrane-induced aggregation, whereas a variant with a low propensity to undergo self-assembly at the membrane (A29E) exhibited no vesicle permeabilization activity (22). Thus, we hypothesized that h-aSyn A53T and m-aSyn Chimera should elicit greater membrane disruption than h-aSyn Chimera and m-aSyn, based on the relative propensities of these variants to undergo membrane-induced aggregation (Figure 7B). To examine this hypothesis, we compared the human, mouse, and chimeric aSyn variants in terms of their ability to disrupt vesicles using a dye release assay. PG-PC SUVs pre-loaded with the fluorescent dye calcein at a concentration leading to self-quenching were incubated with the aSyn variants in monomer form at 37 °C. Membrane permeabilization was assessed by monitoring the increase in fluorescence intensity resulting from dequenching of the dye after its release from the vesicles and dilution in the surrounding buffer. Upon extended incubation, all of the aSyn-SUV mixtures displayed a time-dependent increase in fluorescence, in contrast to the control sample consisting of vesicles without protein (Figure 7C). After 120 h, the fluorescence emission observed for h-aSyn A53T and m-aSyn Chimera (corresponding to a yield of ∼55% of the maximal signal obtained by lysing the vesicles with detergent) was 2.5- and 1.5-fold greater than that observed for h-aSyn Chimera and m-aSyn, respectively (Figure 7C). H-aSyn WT produced a similar increase in fluorescence compared to h-aSyn A53T and m-aSyn Chimera. Together, these data suggested that the human-to-mouse substitutions at positions 121 and 122 interfere with aSyn-mediated vesicle disruption under conditions compatible with aSyn aggregation at the membrane surface.

## Discussion

Understanding how human and mouse aSyn differ in terms of pathobiology is a high priority in the synucleinopathy field not only because of the A/T mismatch at position 53 relevant to familial PD, but also because of the widespread use of genetically engineered mice to investigate molecular underpinnings of aSyn neuropathology *in vivo*. Previously, it was reported that transgenic mice expressing m-aSyn downstream of the Thy-1 promoter show evidence at postmortem examination of extranigral aSyn accumulations similar to those observed in mice expressing WT or mutant forms of h-aSyn (41). Moreover, Lashuel and colleagues (42) showed that endogenous m-aSyn interferes with the formation of Lewy-like inclusions by h-aSyn WT in cultured neurons and in mouse brain. Here we report that m-aSyn has a reduced propensity to elicit dopaminergic cell death compared to h-aSyn A53T in primary midbrain cultures and exhibits a similar effect when expressed from an AAV vector in rat SN. Furthermore, we show for the first time that this differential toxicity is associated with differences in aSyn-mediated vesicle disruption and aggregation at the membrane surface, rather than differences in aSyn fibrillization previously suggested to be important for modulating its toxicity (43). Most importantly, we provide evidence that these characteristics of the overtly toxic h-aSyn A53T variant can be blunted, down to the same level of m-aSyn, by modification of only 2 amino acid residues in positions 121 and 122 of the protein.

Our findings are consistent with the idea that m-aSyn has a reduced propensity to elicit neurotoxicity compared to h-aSyn A53T as a result of attenuation of the toxic effects of threonine 53 in the mouse protein by variations in other sites of the protein. Moreover, these results support the hypothesis that mice show no signs of neurodegeneration despite expressing an aSyn variant with a threonine residue at position 53 at least in part because m-aSyn is intrinsically less toxic than h-aSyn A53T. The lack of nigral degeneration in mouse models with altered expression of proteins that interact functionally with aSyn, including transgenic mice over-expressing LRRK2 (44) and DJ-1 knockout mice (45), could also potentially be explained by the reduced neurotoxicity of endogenous m-aSyn. Nevertheless, despite its lower intrinsic toxicity, m-aSyn can also contribute to dopaminergic cell death in other models relevant to nigral degeneration, including mice treated with MPTP (46). In addition to the intrinsic biochemical differences between h-aSyn A53T and m-aSyn reported here, other factors including differences in neurochemistry between the nigrostriatal systems of mice and humans (47, 48) could also account at least in part for the apparent absence of early-onset aSyn neuropathology in mice in contrast to human A53T patients.

Our data are in line with previous results showing that rat aSyn, identical to m-aSyn except for a single G-to-S mismatch at position 121, triggers less extensive nigral degeneration compared to h-aSyn WT, h-aSyn A30P, or h-aSyn A53T when expressed from a lentiviral vector in rat midbrain (28). Moreover, we observed that rat aSyn has a similar blunted response of nigral dopaminergic cell death compared to mouse aSyn in the rAAV-mediated overexpression model in the rat (29). Another group reported evidence of TH^+^ neuron loss and an accumulation of Lewy-like inclusions in the SN of rats injected intrastriatally with preformed fibrils (PFFs) prepared from recombinant m-aSyn (49). A similar but less pronounced neuropathological phenotype was observed upon injecting a sample of non-fibrillized m-aSyn in rat midbrain, although the interpretation of the results was complicated by uncertainty about the degree of aSyn oligomerization in the injected, non-fibrillar protein sample.

A key finding of this study was the observation in both primary mesencephalic cultures and rat midbrain that (i) h-aSyn Chimera, a variant of h-aSyn A53T with m-aSyn residues introduced at positions 121 and 122, was less neurotoxic than h-aSyn A53T; and (ii) m-aSyn Chimera, a variant of m-aSyn with human aSyn residues introduced at positions 121 and 122, was more neurotoxic than the mouse protein. From these results, we infer that the presence of glycine and serine residues at positions 121 and 122 of m-aSyn (respectively) is largely responsible for offsetting the expected neurotoxic effects of threonine 53 in the murine sequence. In turn, these findings reveal for the first time that residues 121 and 122 play a key role in modulating aSyn-mediated neurodegeneration. Notably, in contrast to the striking effects of human/mouse substitutions at positions 121 and 122, the S87N or N87S substitution had no impact on the ability of human, mouse, or chimeric aSyn variants to trigger dopaminergic cell death in primary midbrain cultures, suggesting that the variations in the two C-terminal residues are unique in their impact on the toxicity. In support of these findings, a number of groups have shown that modifications in the C-terminal domain, including truncation (31, 34) and Y125 phosphorylation (32), affect aSyn neurotoxicity in cellular and animal models.

Previously, m-aSyn was shown to form amyloid-like fibrils with a reduced lag time and a greater elongation rate compared to h-aSyn WT or h-aSyn A53T (25, 26). Here we confirmed these earlier findings and found that two chimeric variants, h-aSyn Chimera and m-aSyn Chimera, undergo less rapid fibrillization than the mouse protein. These results indicate that the human-to-mouse substitutions at positions 121 and 122 are necessary but not sufficient for the enhanced fibrillization rate of m-aSyn relative to h-aSyn A53T. Because the human-to-mouse and mouse-to-human substitutions at positions 121 and 122 do not have opposite effects on the rate of aSyn fibrillization, we infer that the impact of each of these substitutions on aSyn self-assembly is dependent on the sequence context, potentially because residues 121 and 122 are involved in long-range intramolecular interactions that influence the rate of aSyn self-assembly (8, 9). Additional data revealed that variants of h-aSyn A53T and h-aSyn Chimera with the S87N substitution form fibrils with a decreased lag time, consistent with previously reported effects of this mutation (25), whereas variants of m-aSyn and m-aSyn Chimera with the N87S substitution form fibrils with an increased lag time.

Importantly, however, differences in fibrillization rates among the human, mouse, and chimeric aSyn variants correlate only weakly with differences in neurotoxicity. For example, h-aSyn A53T, h-aSyn Chimera, m-aSyn, and m-aSyn Chimera exhibit the following order of fibrillization rates:

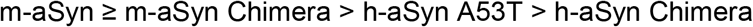

In contrast, the same variants exhibit the following order of neurotoxicity:

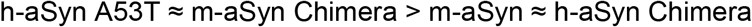

Moreover, despite causing changes in aSyn fibrillization lag times, the S87N and N87S substitutions have no impact on aSyn neurotoxicity in primary midbrain cultures, further highlighting the weak correlation between fibrillization propensity and aSyn-mediated neurodegeneration revealed by our study. The lack of a tight correlation may reflect the fact that in mammalian cells non-membrane-bound aSyn adopts a compact disordered conformation that is apparently incompatible with efficient fibrillization (5). Thus, a perturbation that favors or hinders fibril formation by recombinant, natively unfolded aSyn in solution may have a very different effect on the protein’s conversion to neurotoxic species in its native intracellular environment.

AFM and TEM analyses revealed that human, mouse, and chimeric aSyn formed amyloid-like fibrils with different morphologies under the experimental conditions used here. Namely, m-aSyn and h-aSyn Chimera, the two variants with reduced neurotoxicity, formed both twisted and non-twisted fibrils, whereas only non-twisted fibrils were formed by h-aSyn A53T and m-aSyn Chimera. These results are consistent with solid-state NMR data showing that fibrils formed by h-aSyn WT and m-aSyn differ in terms of protofilament substructure (50, 51), although previous EM analyses revealed no major morphological differences between fibrils prepared from human A53T and mouse aSyn (25, 52). Multiple groups have shown that injecting PFFs into the brains of WT or transgenic rodents initiates the aggregation of endogenous aSyn via a seeding mechanism, leading to the formation of Lewy-like inclusions (49, 53, 54). Moreover, evidence suggests that different types of fibrils (e.g. cylindrical fibrils versus ribbons distinguishable by EM) can act as prion-like ‘strains’ with different abilities to undergo cell-to-cell transmission throughout the brain (37). Our AFM and TEM data imply that m-aSyn and h-aSyn Chimera could potentially form different fibrillar strains (characterized by increased amounts of twisted fibrils) compared to the other aSyn variants, and, therefore, the differences in *in vivo* neurotoxicity reported here could be due to differences in the variants’ abilities to seed aSyn neuropathology in rat brain. However, this interpretation is challenged by evidence that (i) h-aSyn A53T can form twisted fibrils under certain incubation conditions (55); and (ii) PFFs formed from recombinant m-aSyn induce inclusion formation and dopaminergic cell death with greater or similar efficiency (respectively) compared to h-aSyn A53T PFFs following their injection in mouse SN (56), in contrast to the reduced neurotoxicity of m-aSyn compared to h-aSyn A53T reported here. Additional studies are needed to assess whether human, mouse, or chimeric aSyn variants produce different fibrillar strains with different levels of neurotoxicity *in vivo*.

Previously we reported that two familial substitutions, A30P and G51D, increase aSyn neurotoxicity by enhancing the protein’s ability to form aggregates at membrane surfaces (16). Here we show that two aSyn variants with a higher propensity to elicit dopaminergic cell death in primary culture and *in vivo*, h-aSyn A53T and m-aSyn Chimera, undergo membrane-induced self-assembly with greater efficiency compared to two variants with lower neurotoxicity, h-aSyn Chimera and m-aSyn. These results suggest that mouse-to-human substitutions at positions 121 and 122 cause an increase in aSyn neurotoxicity by promoting Syn aggregation at membrane surfaces, whereas human-to-mouse substitutions have the opposite effect. Because all of the aSyn variants examined in this study exhibited similar affinities for PG:PC SUVs, we infer that human-to-mouse substitutions interfere with lipid-induced aSyn aggregation not by favoring binding of the central hydrophobic domain to the membrane, but rather by inducing a conformational change in the C-terminal region that disrupts aSyn self-assembly, potentially via perturbation of long-range interactions (8, 9, 40). Consistent with this scenario, data from several studies suggest that modifications in the C-terminal region (namely, Y125 phosphorylation or nitration, or the nitration of all three C-terminal tyrosine residues) can modulate the conformation of the N-terminal and central hydrophobic domains of the membrane-bound form of the protein (40, 57, 58). These findings support the idea that long-range interactions can occur in the membrane-associated form of aSyn, in turn suggesting that the human-to-mouse substitutions at positions 121 and 122 could potentially influence membrane-induced aSyn aggregation by altering these intramolecular contacts. Moreover, Burré et al. (59) have proposed a model wherein oligomers formed by aSyn at the membrane surface consist of neighboring aSyn molecules with their C-terminal alpha-helices arranged in an anti-parallel configuration. In this arrangement, the aspartate residue at position 121 of human aSyn could potentially participate in electrostatic interactions with positively charged residues on a neighboring subunit, and in turn these interactions could favor the formation of pathological aSyn aggregates at the membrane surface. This idea is also supported by computational data from our group suggesting that intermolecular interactions between the C-terminal domain and central hydrophobic region play a key role in aSyn dimerization at the surface of anionic phospholipid membranes (60).

In the current study and in our earlier work (16, 22), SDS-resistant aSyn oligomers were detected in the lipid fraction of protein/SUV mixtures isolated by lipid flotation. These data indicate that membrane-bound aSyn oligomers were generated upon incubating the protein with SUVs. The formation of these membrane-bound oligomers could occur as a result of (i) aSyn aggregation at the membrane surface, or (ii) aSyn aggregation in solution, followed by the binding of the aggregates to the membrane. Three observations argue against the second scenario. First, we find that aSyn does not form aggregates when incubated in the absence of SUVs under the same conditions as those used in the membrane-induced aggregation assay. Second, the relative rates at which aSyn variants form oligomers in the presence of SUVs correlate poorly with the relative rates at which the variants undergo self-assembly in the absence of phospholipids (16). Third, we recently showed that ENSA, a protein that interacts selectively with membrane-bound aSyn (61), inhibits aSyn aggregation in the presence of SUVs, whereas it does not interfere with aSyn fibrillization in the absence of lipids (22). The fact that ENSA does not inhibit aSyn fibril formation in the absence of SUVs suggests that the inhibitory effect of ENSA on aSyn self-assembly in the presence of lipids must arise from its interaction with aSyn at the membrane surface. In turn, this evidence supports the idea that aSyn forms oligomers on the membrane when incubated with SUVs under the conditions described here.

Importantly, our data reveal a strong correlation between aggregation propensity at membrane surfaces and neurotoxicity among the human, mouse, and chimeric aSyn variants, and thus they further support the hypothesis that lipid-induced self-assembly plays a key role in aSyn-mediated neurodegeneration (11, 16, 18, 22). The importance of membrane-induced aSyn aggregation to the pathogenesis of synucleinopathy disorders is further underscored by evidence that non-membrane-associated aSyn adopts a compact, aggregation-resistant conformation, with little exposure of the central hydrophobic region, in mammalian cells (5). Although it is uncertain why all of the aSyn variants examined here formed PK-resistant aggregates in rat striatum, one explanation could be that the inclusions consist of fibrils derived from both membrane-bound and non-membrane-bound forms of aSyn (20, 21). Accordingly, a decrease in the levels of fibrils formed in the presence of lipids could be offset by an increase in the levels of fibrils formed away from the membrane in the case of m-aSyn, whereas the opposite scenario could be true in the case of h-aSyn A53T. Consistent with this idea, data reported by multiple groups imply that aSyn can undergo various forms of self-assembly, leading to the formation of different types of aggregated species with different degrees of neurotoxicity, in cell culture models, aSyn transgenic mice, and the brains of synucleinopathy patients (62–66). Although our results highlight the importance of membrane-induced aSyn aggregation to aSyn-mediated dopaminergic cell death, we suggest that aSyn aggregates formed both at membrane surfaces and in solution likely contribute to the overall neuropathology of synucleinopathy disorders. Assessing whether differences in the rates or extents of aSyn oligomerization in the absence of membranes also account at least in part for differences in neurotoxicity among the aSyn variants is an important goal for future studies.

A final key outcome of this study was our finding that the relatively neurotoxic variants h-aSyn A53T and m-aSyn Chimera elicit the disruption of synthetic vesicles under conditions favoring membrane-induced aSyn aggregation with greater efficiency than the less toxic variants h-aSyn Chimera and m-aSyn. These data suggest that mouse-to-human substitutions at positions 121 and 122 enhance aSyn neurotoxicity by promoting aSyn-mediated membrane permeabilization, whereas human-to-mouse substitutions have the opposite effect. Unexpectedly, we found that h-aSyn WT triggered vesicle disruption to a similar extent as h-aSyn A53T and m-aSyn Chimera. This observation was inconsistent with the fact that h-aSyn WT exhibited a non-significant trend towards less extensive membrane-induced self-assembly, or with evidence from our earlier studies that h-aSyn WT is considerably less toxic than h-aSyn A53T in primary midbrain cultures (16) and causes neurodegeneration less rapidly than h-aSyn A53T in rat midbrain (67). As one possibility, the current findings can be interpreted to mean that h-aSyn WT engages in different molecular interactions during aggregation at the membrane surface compared to h-aSyn A53T or m-aSyn Chimera, resulting in an increase in the extent of membrane permeabilization relative to the total amount of aggregate formed. Consistent with this idea, evidence from multiple studies suggests that h-aSyn WT and familial aSyn mutants can form different types of amyloid-like fibrils (68–70). Overall, our data reveal a strong correlation between vesicle disruption and neurotoxicity among the human, mouse, and chimeric aSyn variants, and thus they support the notion that membrane permeabilization associated with lipid-induced self-assembly plays a key role in aSyn-mediated neurodegeneration. These findings are consistent with evidence that h-aSyn E57K, a variant with a high propensity to form aggregates on a supported lipid bilayer and elicit membrane damage, causes extensive dopaminergic cell death when expressed in rat SN (19, 35).

Previous findings reported by Volles et al. (71, 72) suggest that aSyn oligomers formed in the absence of membranes fail to permeabilize low-diameter vesicles composed of a mixture of anionic and zwitterionic lipids. Accordingly, we infer that the vesicle disruption observed in aSyn-SUV mixtures in this study and in our earlier work (22) results from aSyn self-assembly at the membrane surface. These observations further corroborate the idea that aSyn forms oligomers on the membrane when incubated with SUVs under the conditions described in this study. Moreover, oligomeric m-aSyn prepared in the absence of membranes was reported by Volles et al. (71) to permeabilize vesicles composed of anionic lipids with a similar efficiency compared to oligomeric h-aSyn A53T, contrary to the difference in the variants’ neurotoxicity reported here. This observation suggests that vesicle disruption associated with lipid-induced aSyn aggregation plays a more important role than membrane permeabilization by preformed oligomers in aSyn-mediated neurodegeneration *in vivo*.

### Conclusions

We report for the first time that m-aSyn has a reduced ability to elicit dopaminergic cell death compared to h-aSyn A53T in primary mesencephalic cultures and in rat midbrain. Our results highlight the central importance of membrane-induced aSyn aggregation and vesicle disruption in aSyn neurotoxicity, and they reveal a C-terminal sequence motif encompassing residues 121 and 122 with an important role in modulating these activities. Collectively, these findings provide a strong rationale for developing therapies aimed at inhibiting aSyn aggregation at membrane surfaces, including interventions that target the C-terminal domain, as a strategy to slow neurodegeneration in the brains of synucleinopathy patients (16, 18, 22).

## Experimental Procedures

### Materials

Unless otherwise stated all chemicals were purchased from Sigma-Aldrich (St. Louis, MO or Stockholm, Sweden). DMEM, FBS, penicillin-streptomycin, Lipofectamine 2000, Optimem, Trypsin-EDTA, ProLong Gold Antifade Reagent with DAPI, Proteinase K (25530–049), and the Virapower Adenoviral Expression System were obtained from Invitrogen (Carlsbad, CA). RPMI medium, phenol red-free RPMI medium, Nuserum, Plus slides (J3800AMNZ), Phosphatase Inhibitor Mini Tablets, bicinchoninic acid (BCA) assay reagents, Coomassie Brilliant Blue, and the Immobilon-FL PVDF membrane was purchased from Thermo Fisher Scientific (Rockford, IL). The QuickTiter Adenovirus Titer Elisa kit was purchased from Cell Biolabs (San Diego, CA). Antisedan, Temgesic, and the Fentanyl/Dormitor mixture were obtained from Apoteksbolaget, (Stockholm, Sweden). The Vectastain ABC kit was purchased from Vector Laboratories, Inc. (Burlingame, CA), and 3,3’-diaminobenzidine (DAB Safe) was obtained from Saveen Werner (Limhamnsvägen, Sweden). Isopropyl β-D-1-thiogalactopyranoside, ampicillin, and PMSF were purchased from Gold Biotechnology (St. Louis, MO). Recombinant Syk, Src, and Fyn kinases were obtained from SignalChem (Richmond, BC, Canada). L-α-phosphatidylcholine (egg PC), L-α-phosphatidylglycerol (egg PG), and phospholipid extrusion membranes were purchased from Avanti Polar Lipids (Alabaster, AL). Iodixanol and the enhanced chemifluorescence (ECF) substrate were obtained from GE Life Sciences (Pittsburgh, PA). The nitrocellulose membrane was purchased from Bio-Rad Laboratories (Hercules, CA). 100 kDa spin filters and 10 kDa concentration filters were obtained from Millipore (Billerica, MA or Solna, Sweden). SH-SY5Y cells were purchased from ATCC (Manassas, VA) (catalog # CRL-2266, RRID:CVCL_0019), and the PC12 Tet-Off cell line (RRID:CVCL_V361) was obtained from Clontech Laboratories (Mountain View, CA). Cell line authentication and mycoplasma testing were carried out by IDEXX BioResearch (Columbia, MO) (cell line identities were confirmed via species-specific PCR evaluation and STR profiling, and both cell lines were shown to be mycoplasma negative). A plasmid encoding murine Syk fused at its C-terminus with EGFP (Syk-GFP) (73) was provided by Dr. Robert Geahlen (Purdue University), and vectors encoding human GST-Src and GST-Fyn (74) were provided by Dr. Laurie Parker (University of Minnesota), with permission from Dr. Benjamin Turk (Yale University).

### Antibodies

The following antibodies were used in these studies: Primary cell culture – chicken anti-MAP2 (CPCA-MAP2, RRID:AB_2138173, EnCor Biotechnology, Gainesville, FL); rabbit anti-tyrosine hydroxylase (TH) (AB152, RRID:AB_390204, Millipore, Bellerica, MA); anti-rabbit IgG-Alexa Fluor 488 (R37116, RRID:AB_2556544) and anti-chicken IgG-Alexa Fluor 594 (A11042, RRID:AB_142803) (Thermo Fisher Scientific, Rockford, IL).

*In vivo* experiments – human-specific syn211 (36-008, RRID:AB_310817, Millipore, Germany); mouse anti-aSyn, clone 42 (Syn-1) (610787, RRID:AB_398108, BD Biosciences, UK); rabbit anti-TH (P40101-0, RRID:AB_461064, Pel-freeze, USA); rabbit anti-VMAT2 (ab81855, RRID:AB_2188123, Abcam, UK or 20042, RRID:AB_10013884, Immunostar, USA); mouse anti-HuC/HuD, clone 16A11 (A21271, RRID:AB_221448, Thermo Scientific, USA); biotinylated secondary antibodies (Vector Laboratories Inc., USA).

Western blotting – mouse anti-aSyn, clone 42 (Syn-1) (610787, RRID:AB_398108, BD Biosciences, San Jose, CA); mouse anti-phosphotyrosine, clone 27B10 (APY03-HRP, Cytoskeleton, Denver, CO); rabbit anti-pY125-aSyn (ab131466, RRID:AB_11158765, Abcam, Cambridge, MA, or A1219, BioVision, Milpitas, CA); mouse anti-β-actin, clone AC-74 (A5316, RRID:AB_476743, Sigma-Aldrich); mouse anti-GFP, clone B-2 (sc-9996, RRID:AB_627695, Santa Cruz Biotechnology, Dallas, TX); rabbit anti-Src (2108, RRID:AB_331137), rabbit anti-Fyn (4023, RRID:AB_2108441), AP-linked anti-mouse IgG (7056, RRID:AB_330921), and AP-linked anti-rabbit IgG (7054, RRID:AB_2099235) (Cell Signaling Technology, Danvers, MA); goat anti-mouse IR-Dye 680RD (92568070, RRID:AB_2651128) and goat anti-rabbit IR-Dye 800CW (92532211, RRID:AB_2651127) (Li-COR, Lincoln, NE).

### Preparation of bacterial expression constructs

Bacterial expression constructs encoding h-aSyn WT (pT7-7-h-aSyn), h-aSyn A53T (pT7-7-h-aSyn-A53T), and m-aSyn (pT7-7-m-aSyn) were described previously (26). A construct encoding h-aSyn Chimera was generated by replacing the BamHI-HindIII fragment of pT7-7-h-aSyn-A53T (encompassing base pairs 354-423 of the h-aSyn A53T cDNA) with the equivalent fragment from pT7-7-m-aSyn. A construct encoding m-aSyn Chimera was generated by replacing the BamHI-HindIII fragment of pT7-7-m-aSyn (encompassing base pairs 354-423 of the m-aSyn cDNA) with the equivalent fragment from pT7-7-h-aSyn-A53T. pT7-7 constructs encoding A53T/D121G, A53T/N122S, A53T/S87N, h-aSyn Chimera S87N, m-aSyn N87S, and m-aSyn Chimera N87S were generated via site-directed mutagenesis using the QuikChange method (Stratagene). The sequence of the DNA insert in each bacterial expression construct was verified using an Applied Biosystems (ABI3700) DNA sequencer (Purdue Genomics Core Facility).

### Adenoviral vector design and virus production

Adenoviral constructs encoding h-aSyn WT and h-aSyn A53T were described previously (16, 75). cDNAs encoding h-aSyn Chimera, A53T/D121G, A53T/N122S, A53T/S87N, and h-aSyn Chimera S87N (obtained using the pT7-7 constructs outlined above as PCR templates) were subcloned as KpnI-XhoI fragments into the entry vector pENTR1A. cDNAs encoding m-aSyn, m-aSyn Chimera, and m-aSyn Chimera N87S were subcloned as SalI-XhoI fragments into pENTR1A. Inserts from the pENTR1A constructs were then transferred into the pAd/CMV/V5 adenoviral expression vector via recombination using Gateway LR Clonase. The sequence of the DNA insert in each adenoviral construct was verified using an Applied Biosystems (ABI3700) DNA sequencer (Purdue Genomics Core Facility). Adenoviral constructs were packaged into virus via lipid-mediated transient transfection of the HEK 293A packaging cell line. Purified adenoviral particles were titered using the QuickTiter Adenovirus Titer Elisa kit.

### AAV vector design and virus production

Adeno-associated viral vectors of serotype 5 (AAV5) were comprised of a genome cassette flanked by inverted terminal repeats (ITR2). The cassette contained a chicken beta-actin promoter in combination with a CMV enhancer element (CBA) leading to expression of one of the four different transgenes. Genes were PCR-amplified with the plasmids pENTR1A h-aSyn A53T, pENTR1A h-aSyn Chimera, pENTR1A m-aSyn, and pENTR1A m-aSyn Chimera as templates and subcloned as HindIII-SpeI fragments in the pTR-UF20 plasmid. The validity of the constructs was confirmed by restriction analysis and Sanger sequencing. Downstream of each cDNA insert was an h-SV40 polyA transcription termination sequence. HEK293 cells were co-transfected at a confluency of 70-80% using the calcium-phosphate precipitation method. Plasmids used here encode essential adenoviral packaging and AAV5 capsid genes as previously described (76). Three days after transfection, cells were harvested in PBS and lysed by performing three freeze-thaw cycles in a dry ice/ethanol bath. The lysate was then treated with benzonase and purified using a discontinuous iodixanol gradient followed by Sepharose Q column chromatography (77). Vectors were concentrated using a 100 kDa molecular weight cut-off column, and titers of the stock solution were determined by qPCR using primers and probes targeting the ITR sequence. Before being used in an experiment, vectors were diluted in PBS, pH 7.4 and re-titered, yielding the values reported in the Results and in Figure 3.

### Preparation and treatment of primary mesencephalic cultures

Primary midbrain cultures were prepared via dissection of day 17 embryos obtained from pregnant Sprague-Dawley rats (Harlan, Indianapolis, IN, USA) as described (16, 75). All of the procedures involving animal handling were approved by the Purdue Animal Care and Use Committee. The cells were plated on poly-L-lysine-treated 48-well plates at a density of 163,500 cells per well. Five days after plating, the cells were treated with cytosine arabinofuranoside (AraC) (20 µM, 48 h) to inhibit the growth of glial cells. At this stage (i.e., 7 days in vitro (DIV)), the neurons appeared differentiated with extended processes. The cultures were transduced with adenoviruses encoding the aSyn variants for 72 h at a multiplicity of infection (MOI) ranging from 7 to 15 to ensure equal expression levels as determined via Western blotting. After incubation for an additional 24 h in fresh medium without virus, the cells were fixed in 4% (w/v) PFA in PBS (pH 7.4). The cells were then washed in PBS and incubated with chicken anti-MAP2 (1:2000) and rabbit anti-tyrosine hydroxylase (TH) 1:500) for 24 h. After an additional wash in PBS, the cultures were incubated with AlexaFluor 594-conjugated goat anti-chicken and AlexaFluor 488-conjugated goat anti-rabbit secondary antibodies (1:1000) for 1 h at 22 °C. ProLong Gold Antifade Reagent with DAPI was applied to each well, and a coverslip was added.

### Measurement of neuronal viability in midbrain cultures

Relative dopaminergic cell viability was evaluated as described (16, 75). MAP2^+^ and TH^+^ cells were counted in a blinded manner in at least 10 randomly chosen observational fields (approximately 500-1000 neurons total) for each experimental condition using a Nikon TE2000-U inverted fluorescence microscope (Nikon Instruments, Melville, NY) equipped with a 20x objective. The data are expressed as the percentage of MAP2^+^ neurons that are also TH^+^ to correct for variation in cell density. Each experiment was repeated a minimum of 3 times with cultures isolated from different pregnant rats.

### Animals for in vivo experiments

Young adult Sprague-Dawley rats received from Charles River (Kisslegg, Germany) were housed on a 12 h light-dark cycle with *ad libitum* access to water and food. All experimental procedures were approved by the Ethical Committee for Use of Laboratory Animals in the Lund-Malmö region.

### Stereotaxic surgery

Rats were anesthetized via i.p. injection of a 20:1 mixture of Fentanyl and Dormitor (6 mL/kg). After placing each animal into a stereotaxic frame (Stoelting, Wood Dale, USA), the SN was targeted unilaterally using the following coordinates: anteroposterior (AP), −5.0 mm; mediolateral (ML), −2.0 mm from bregma; and dorsoventral (DV), −7.2 mm from the dura. The tooth bar was adjusted to −2.3 mm. A pulled glass capillary (about 60-80 µm in diameter) was attached to a 5 µL Hamilton syringe equipped with a 22s gauge needle and used to deliver 2 µL of rAAV5 vector solution at a pulsed injection of 0.1 µL every 15 sec. After delivery of the vector, the capillary was left in place for 5 min, retracted 0.1 mm, and after 1 min it was slowly removed from the brain. After closing the wound with clips, Antisedan and Temgesic were administered s.c. as an analgesic treatment and to reverse the anesthesia.

### Histology

An overdose of sodium pentobarbital was used to kill rats 8 weeks after vector delivery. Animals were perfused via the ascending aorta first with 50 mL of 0.9% (w/v) NaCl followed by 250 mL of ice-cold 4% (w/v) PFA in 0.1 M phosphate buffer, pH 7.4, for 5 min. Brains were taken out and post-fixed in 4% (w/v) PFA for 24 h and then incubated in 25% (w/v) sucrose for cryoprotection. Brains were cut on a microtome (Microm HM450, Thermo Scientific, USA) in 35 µm-thick coronal sections and placed into antifreeze solution (0.5 M phosphate buffer, 30% (v/v) glycerol, 30% (v/v) ethylene glycol) at −20 °C for long term storage until further processed.

Immunohistochemical staining was performed on free-floating brain sections. Specimens were rinsed with Tris-buffered saline (TBS) buffer (5 mM Tris-HCl, 15 mM NaCl, pH 7.6). Only TH staining required an antigen retrieval step carried out for 30 min at 80 °C using Tris/EDTA buffer (10 mM Tris-HCl, 1 mM EDTA, pH 9.1). This step was terminated by repeated washes in TBS at 22 °C. Endogenous peroxidase activity was quenched using a solution of 3% (v/v) H_2_O_2_ and 10% (v/v) methanol in TBS buffer for 30 min, and the reaction was stopped by washing sections 3 times with TBS. To block unspecific binding sites, sections were incubated for 30 min in 0.05% (v/v) Triton X-100 in TBS buffer (TBS-T) containing 5% (v/v) normal serum from the same species as that in which the secondary antibody was raised. The primary antibodies were then applied in 1% (w/v) BSA in TBS-T overnight at 22 °C: human-specific syn211 at 1:100,000; Syn-1 at 1:10,000; TH at 1:5,000; VMAT2 ab81855 at 1:4,000, or 20042 at 1:10,000; HuC/HuD 1:200. The next day sections were washed in TBS-T, and the appropriate biotinylated secondary antibody (1:200) was applied in 1% (w/v) BSA in TBS-T for 1 h. Specimens were rinsed once more with TBS-T and further incubated in a solution containing avidin-biotin-peroxidase complexes (Vectastain ABC kit, Vector Laboratories Inc., USA) for 1 h. Staining was visualized using 3,3’-diaminobenzidine (DAB Safe) and 0.01% (v/v) H_2_O_2_. Sections were then mounted on chromatin-gelatin coated glass slides, dehydrated in increasing alcohol solutions, cleared with xylene, and applied to coverslips using DPX (06522, Sigma-Aldrich, Sweden).

For cresyl violet staining, sections were mounted on chromatin-gelatin coated glass slides and dried overnight. Sections were then hydrated in decreasing alcohol solutions and stained for 30 sec in 0.5% (w/v) cresyl violet + 0.1% (v/v) acetic acid. After rinsing specimens with H_2_O, sections were dehydrated in increasing alcohol solutions, cleared in xylene, and coverslipped using DPX.

To visualize insoluble aggregates, sections were treated with Proteinase K. For this purpose, specimens were first rinsed with potassium-containing phosphate-buffered saline (KPBS) and then heat-treated at 80°C for 30 min in KPBS. Sections were then quenched using 3% (v/v) H_2_O_2_ and 10% (v/v) methanol in KPBS for 30 min. After mounting sections on plus slides, the sections were incubated in a KPBS solution containing 10 µg/mL Proteinase K for either 5 or 45 min. The same staining protocol was applied as described above using the aSyn-specific primary antibody Syn-1 at 1:1,000.

### Stereological analysis

In order to estimate surviving TH^+^ nigral neurons, an unbiased stereological quantification using the optical fractionator method (78, 79) was applied. The implementation of a random starting position and systematic sampling was done using the Visiopharm VIS software (version 4.5.5.433 Visiopharm A/S, Denmark) and a Nikon Eclipse 80i light microscope fitted with an xy motorized stage, a motorized z-axis (Prior, UK), and a microcator (Heidenheim, Germany). Through the rostro-caudal axis of the SN, every 6^th^ section was stained for TH. Regions of interest were outlined with a 4x objective, and counting was executed using a 60x oil-immersion objective with a numerical aperture of 1.4. Counting parameters were set to reach at least 100 cell counts per hemisphere. Estimation of the total number of TH^+^ cells was performed by a single blinded investigator. The coefficient of error due to the sampling was calculated based on Gundersen and Jensen (80), and values lower than 0.1 were considered acceptable. The contralateral uninjected side of each animal served as an internal control, and values were expressed as a percentage of cell counts on the injected side relative to the intact side (mean ± SEM).

### Densitometry

The optical density of TH^+^ and VMAT2^+^ fibers was measured on digital images of coronal striatal sections using the Zeiss microscope (Axio Zoom.V16, Zeiss, Germany). The striatum of every 24^th^ section in the rostro-caudal axis – in total 6 sections per brain – was outlined using ImageJ, and optical density readings were corrected for non-specific background using density measurements from the *corpus callosum* of each animal. Data consisting of means ± SEM from all brains are expressed as a percentage of contralateral side values.

### Protein purification

Recombinant aSyn variants were purified from BL21 (DE3) cells transformed with pT7-7 constructs as described (16). The final purity of each protein was found to be > 98% as determined by SDS-PAGE with Coomassie Blue staining.

### aSyn fibrillization

Lyophilized aSyn was dissolved in PBS (pH 7.4) with 0.02% [v/v] NaN_3_, and the solution was filtered by successive centrifugation steps through a 0.22 µm spin filter and a 100 kDa centrifugal filter to isolate monomeric protein. The stock solution was dialyzed against the same buffer (24 h at 4 °C) to remove excess salt. Fibrillization solutions of aSyn were prepared by diluting the appropriate stock solutions of dialyzed protein with buffer to a final concentration of 35 µM in the wells of a 96-well plate. To determine the extent of aSyn fibrillization, thioflavin T (ThioT, final concentration, 20 µM) was added to the fibrillization solutions, which were incubated at 37 °C with shaking at the ‘normal’ setting in a Spectra Fluor Plus or Genios plate reader (Tecan, Upsala, Sweden). ThioT fluorescence was measured with excitation at 440 nm and emission at 490 nm. Mean ThioT fluorescence data determined from 3 or 4 technical replicates (defined here as samples treated identically in a single experiment) were normalized using the following equation:

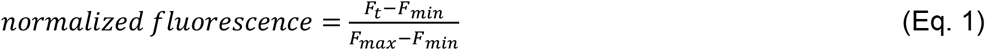

where F_t_ is the ThioT fluorescence emission intensity at time t, and F_min_ and F_max_ are the minimum and maximum fluorescence intensities during the incubation, respectively.

### Lipid vesicle preparation

Small unilamellar vesicles (SUVs) (diameter = 50 nm) were prepared as described (16). We opted to examine SUVs composed of egg PG and egg PC (1:1 mol/mol) because they contain anionic lipids necessary for aSyn membrane interactions (12) and this lipid composition is compatible with producing stable 50 nm vesicles (16). Egg PG and egg PC suspended in chloroform were mixed in a round bottom flask. The chloroform was evaporated under a nitrogen stream and further dried under vacuum for 1 h. Dried lipids were suspended in PBS, and 50 nm SUVs were prepared by extruding the suspension through a Whatman membrane. The size of the vesicles was confirmed by dynamic light scattering. Lipid vesicles were stored at 4 °C prior to use.

### Circular dichroism

Far-UV CD measurements were performed as described (16) using a Chirascan SC spectrometer (Applied Photophysics, Leatherhead, UK). Solutions of recombinant aSyn (5 µM in 20 mM KPi, pH 7.4) in the absence or presence of SUVs (protein-to-lipid ratio, 1:50 to 1:1600, mol/mol) were analyzed in a 1 mm quartz cuvette at 25°C. The ellipticity at 222 nm was recorded and background corrected to eliminate scattering signals arising from the SUVs. The mean residue molar ellipticity at 222 nm ([θ]_*MR*,222_) following equation:

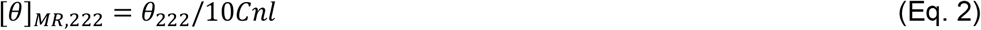

where θ_222_ is the observed (background corrected) ellipticity at 222 nm in millidegrees, C is the protein concentration in M, n is the number of amino acid residues in the protein, and l is the path length of the cuvette in cm. Binding curves were generated by plotting [θ]*_MR_*_,222_ versus the lipid concentration. Curves were analyzed by fitting to the following equation:

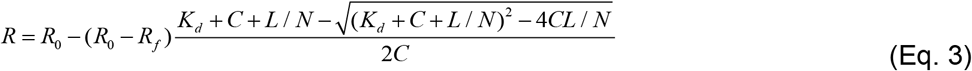

where *R* is the measured [θ]_*MR*,222_ at a given lipid concentration, *R*_0_ is [θ]_*MR*,222_ in the absence of lipid, *R_f_* is [θ]_*MR*,222_ in the presence of saturating lipid, *L* is the total lipid concentration, *C* is the total protein concentration, *Kd* is the apparent macroscopic dissociation equilibrium constant, and *N* is the binding stoichiometry (lipids/protein) (16, 81).

To determine the maximal helical content of each aSyn variant, we used the following equations:

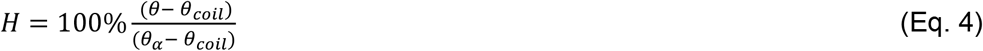

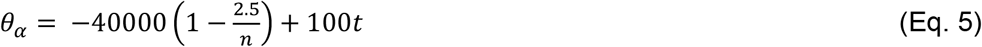

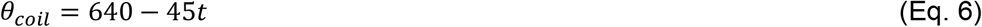

where H is the maximal % helicity, θ is [θ]_*MR*,222_ in the presence of saturating lipid (R*_f_* in Eq. 3), θ_α_ is the [θ]_*MR*,222_ value of an idealized α-helical peptide, θ_coil_ is the [θ]_*MR*,222_ value of an idealized random coil peptide, n is the number of amino acid residues in the protein (= 140), and t is the temperature (25 °C) (16, 82, 83).

### Lipid flotation assay

Lipid flotation analyses were carried out as described (16, 22). Lyophilized aSyn was resuspended in PBS with 0.02% (w/v) NaN_3_, and the solution was dialyzed against the same buffer to remove salts and filtered by successive centrifugation steps through a 0.22 µm spin filter and a 100 kDa centrifugal filter to isolate monomeric protein. aSyn (40 µM) was incubated with PG:PC SUVs (protein-to-lipid ratio, 1:20 mol/mol) at 37°C for 72 h in a total volume of 60 µL. After incubation, the sample was mixed with 4 mL of 30% (v/v) iodixanol solution and overlaid with 7.0 mL of 25% (v/v) iodixanol and 350 µL of 5% (v/v) iodixanol in a polyalomar tube (Beckman, Miami, FL). All of the above iodixanol solutions were prepared in lipid flotation buffer (10 mM HEPES, pH 7.4, 150 mM NaCl). The samples were spun at 200,000 x g in a Beckman SW 41 Ti rotor for 4 h. The membrane fraction was carefully collected from the 5% iodixanol fraction at the top of the gradient, concentrated using a 10 kDa spin filter, and analyzed via Western blotting using a primary antibody specific for aSyn (Syn-1) (1:1,500) (16, 22). In general, we found that the rate of aSyn aggregation increased in the presence of SUVs that had been stored for ∼2-4 weeks at 4 °C (16).

### Synthetic vesicle disruption assay

Calcein-loaded egg PG:PC SUVs were prepared as described (84) with some modifications. A solution of calcein (170 mM) was prepared in H_2_O using NaOH to adjust the pH to 7.4. The final osmolality was 280 mOsm/kg. The calcein solution was used to resuspend dried egg PG:PC lipids, and the suspension was extruded through a Whatman membrane to generate 50 nm SUVs. Calcein-containing vesicles were isolated from free calcein via gel filtration through a Sephadex G-50 column pre-equilibrated with PBS, pH 7.4, 0.02% (w/v) NaN_3_ (280 mOsm/kg). Fractions containing isolated vesicles were pooled and stored at 4 °C until use. Calcein-loaded vesicles were found to be very stable, with no spontaneous dye leakage observed over several weeks.

For membrane disruption experiments, monomeric aSyn variants were isolated as described above (see ‘Lipid Flotation Assay’). Each aSyn variant (40 µM) was incubated with calcein-loaded PG:PC SUVs (protein-to-lipid ratio, 1:20 mol/mol) at 37°C in a total volume of 40µL. At each time point, 3 μL of the reaction mixture was diluted into 180 μL of PBS, and the diluted samples were analyzed with a Fluoromax-3 spectrofluorometer (Horiba Scientific, Edison, New Jersey) (excitation wavelength, 485 nm; emission wavelengths, 505-530 nm; slit width, 1 nm). To determine the maximum dye release, vesicles were lysed by adding 5 µL of 1% (v/v) Triton X-100. The % leakage at time t was determined using the following equation:

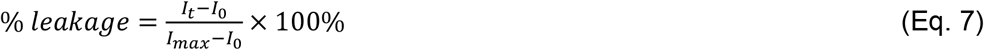

where I_t_ is the fluorescence emission intensity at time t, I_0_ is the intensity at time 0, and I_max_ is the maximal intensity determined after detergent lysis of the vesicles (intensity values were determined at 515 nm).

### AFM analysis of aSyn amyloid-like fibrils

A mica surface functionalized with 1-(3-aminopropyl) silatrane (APS) was used to image aSyn amyloid-like fibrils by AFM (18, 85–87). APS-mica was prepared by incubating freshly cleaved mica in a solution of APS (167 μM) for 30 min and then rinsed with deionized water and dried with an argon stream. A suspension of aSyn fibrils (10 µL, prepared as described above under ‘aSyn fibrillization’ with an incubation time of ∼100 h and diluted 1/20 in deionized water) was deposited onto the APS-mica surface, and the sample was incubated for 2 min, rinsed with deionized water, and dried under an argon stream. The sample was imaged with an AFM Nanoscope VIII system (Bruker, Santa Barbara, CA) using MSNL probes (Cantilever F with spring constant 0.6 N/m), operating in air in peak force mode. Images were acquired over a few randomly selected locations. Images were analyzed using Gwyddion and FemtoScan online software (Advanced Technologies Center, Moscow, Russia) (88, 89).

### TEM analysis of aSyn amyloid-like fibrils

The morphology of aSyn amyloid-like fibrils (prepared as described above under ‘aSyn fibrillization’ with an incubation time of ∼100 h) was analyzed by negative stain biological TEM (90). For biological sample preparation by negative staining, which is a sample preparation technique that imparts the necessary contrast for viewing during biological TEM imaging, 3 µL of aSyn sample solution (35 µM) was pipetted on a discharged carbon-coated copper TEM grid substrate. Subsequently, the sample was washed with deionized water carefully without letting it dry and stained with a 1% (w/v) phosphotungstic acid solution (3 µL), which was left in contact with the protein on the grid for 1 min. The excess solution was then removed by blotting with filter paper, and the sample was imaged using an FEI Tecnai G2 20 Transmission Electron Microscope operating at 200 KV.

### Sample size estimation

Samples sizes for studies of primary neuron viability, membrane-induced aSyn aggregation, and aSyn-mediated membrane permeabilization were estimated via power analysis based on our previous studies (16, 22). Power analyses were designed to determine the number of biological replicates required to detect a ∼20% or ∼30% difference from control, with α = 0.05 (type I error) and power = 0.80 (statistical power). Biological replicates are defined here as samples treated identically in independent experiments carried out on different days. For *in vivo* experiments, sample sizes were estimated based on previous data collected by our group and others for the A53T variant.

### Statistical analysis

Statistical analyses were carried out using GraphPad Prism 6.0 (La Jolla, CA). Primary neuron viability data, in vivo stereology and densitometry data, densitometry data from Western blots, and calcein dye release data were analyzed via ANOVA followed by Tukey’s multiple comparisons *post hoc* test for normally distributed measurements. In analyzing percentage dye release data and percentage cell viability data by ANOVA, square root transformations were carried out to conform to ANOVA assumptions. Normalized FFN-102 fluorescence data were subjected to a log transformation to account for skewness in the data. The log-transformed data were analyzed using an approach that accounts for comparison across experiments conducted on different days. Log-transformed fluorescence emission values for multiple treatment groups were compared using a general linear model implemented in the GLM procedure of SAS Version 9.3 followed by Tukey’s multiple comparisons *post hoc* test (Cary, NC). Kruskal-Wallis followed by a Dunn’s multiple comparisons test or a two-tailed Mann-Whitney test were used as non-parametric tests. Differences were considered significant for p<0.05. Unless otherwise stated, values of n indicated in the Figure Legends refer to the number of independent biological replicates.

## Acknowledgments

We thank Björn Anzelius, Anneli Josefsson, Ulla Samuelsson, and Ulrika Sparrhult-Björk (Lund University) for excellent technical assistance and Aswathy Chandran (Purdue University) for valuable discussions. We are also grateful to Dr. Christopher Gilpin (Purdue University) for advice on TEM analyses.

