## Supplemental Information for "Two C-terminal sequence variations determine differential neurotoxicity between human and mouse α-synuclein"

#### SI Figure Legends

**Figure S1. Differences in antibody recognition of the different aSyn variants. Related to Figure 3.** Adjacent rat midbrain sections of high-titer groups were stained with human aSyn-specific syn211 antibody (A-E) or for comparison with pan-aSyn-specific Syn-1 antibody (F-J). Corresponding contralateral sides are shown in panels A and F, respectively.

**Figure S2. Human/mouse mismatches at positions 121 and 122 do not affect the ability of aSyn to form Proteinase K-resistant aggregates in rat striatum. Related to Figure 4.** Striatal specimens of animals from the highest vector group were incubated in the presence of Proteinase K for 0 min (B-E), 5 min (G-J), or 45 min (L-O) and stained for aSyn to reveal the formation of digestion-resistant aggregates. The digestion of endogenous (and therefore soluble) aSyn was monitored in the CA2/CA3 region of the hippocampus (A, F, K). Scale bar, 25 µm.

**Figure S3. Effects of human/mouse substitutions at position 87 on fibrillization rates of aSyn variants. Related to Figure 5.** The formation of amyloid-like fibrils was monitored in solutions of h-aSyn A53T and h-aSyn A53T/S87N (A), h-aSyn Chimera and h-aSyn Chimera S87N (B), m-aSyn and m-aSyn N87S (C), or m-aSyn Chimera and m-aSyn Chimera N87S (D) (35 µM of each). The protein solutions were incubated at 37°C with constant agitation and analyzed at various times for thioflavin T fluorescence. The graphs show the mean normalized fluorescence (determined from 3 or 4 technical replicates in each experiment) plotted against the incubation time.

### SI Tables

**Table S1. Membrane affinities and maximum helix content of aSyn variants titrated with PG:PC SUVs[a]**

| Variant | $K_d$[b] (μM) | ellipticity minimum (x $10^3$ deg·cm$^2$/dmol) | maximum helicity (%) |
|---|---|---|---|
| h-aSyn-WT | 1.5 ± 0.4 | -18.6 ± 0.8 | 50 ± 3 |
| h-aSyn-A53T | 2.5 ± 0.3 | -20.5 ± 0.4 | 55 ± 2 |
| m-aSyn | 2.4 ± 0.2 | -20.4 ± 0.4 | 55 ± 2 |
| h-aSyn-Chimera | 2.6 ± 0.2 | -21.1 ± 0.4 | 56 ± 2 |
| m-aSyn-Chimera | 1.6 ± 0.2 | -17.7 ± 0.3 | 47 ± 2 |

[a]Values (± standard error) were determined from the far-UV CD data in Fig. 11A using equations 3-6; the concentration of aSyn was 5 μM.
[b]The CD data obtained for each variant were fit to equation 3 with the *N* value (binding stoichiometry) set to 160, the value obtained by fitting the h-aSyn-WT data to equation 3 without any constraints on *N*.

**Table S2. *P* values for vesicle permeabilization data in Figure 7C[1]**

| Comparison | Time (h) | | | | |
|---|---|---|---|---|---|
| | 24 | 48 | 72 | 96 | 120 |
| ctrl vs. h-aSyn | **** | **** | **** | **** | **** |
| ctrl vs. h-aSyn A53T | **** | **** | **** | **** | **** |
| ctrl vs. m-aSyn Chimera | **** | **** | **** | **** | **** |
| ctrl vs. h-aSyn Chimera | **** | **** | **** | **** | **** |
| ctrl vs. m-syn | **** | **** | **** | **** | **** |
| h-aSyn vs. h-aSyn A53T | ns | ns | ns | ns | ns |
| h-aSyn vs. m-aSyn Chimera | ns | ns | ns | ns | ns |
| h-aSyn vs. h-aSyn Chimera | ns | ns | **** | **** | **** |
| h-aSyn vs. m-aSyn | ns | ns | ** | ** | **** |
| h-aSyn A53T vs. m-aSyn Chimera | ns | ns | ns | ns | ns |
| h-aSyn A53T vs. h-aSyn Chimera | ns | *** | **** | **** | **** |
| h-aSyn A53T vs. m-aSyn | ns | ** | ** | * | **** |
| m-aSyn Chimera vs. h-aSyn Chimera | ns | * | *** | **** | **** |
| m-aSyn Chimera vs. m-aSyn | ns | ns | ns | ns | *** |
| h-aSyn Chimera vs. m-aSyn | ns | ns | ns | *** | * |

[1]Two-way ANOVA; *p<0.05, **p<0.01, ***p<0.001, ****p<0.0001.

Figure S1.

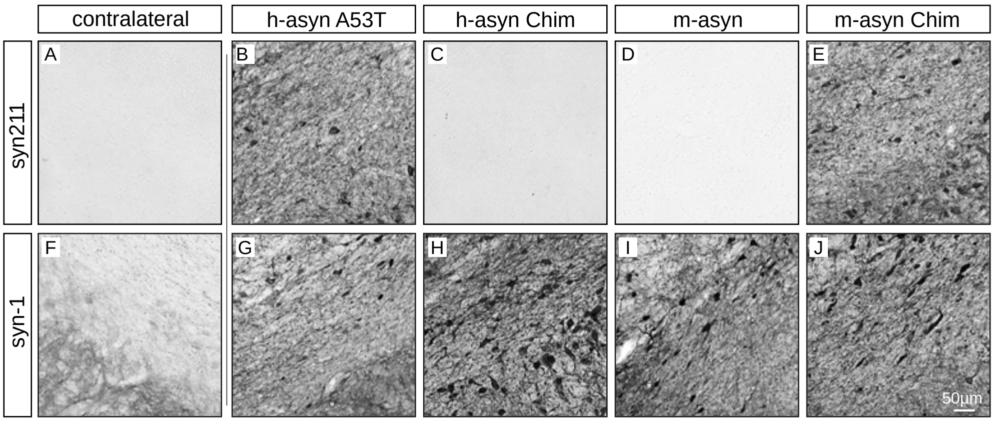

contralateral
h-asyn A53T
h-asyn Chim
m-asyn
m-asyn Chim
syn211
syn-1
A
B
C
D
E
F
G
H
I
J
50µm

Figure S2

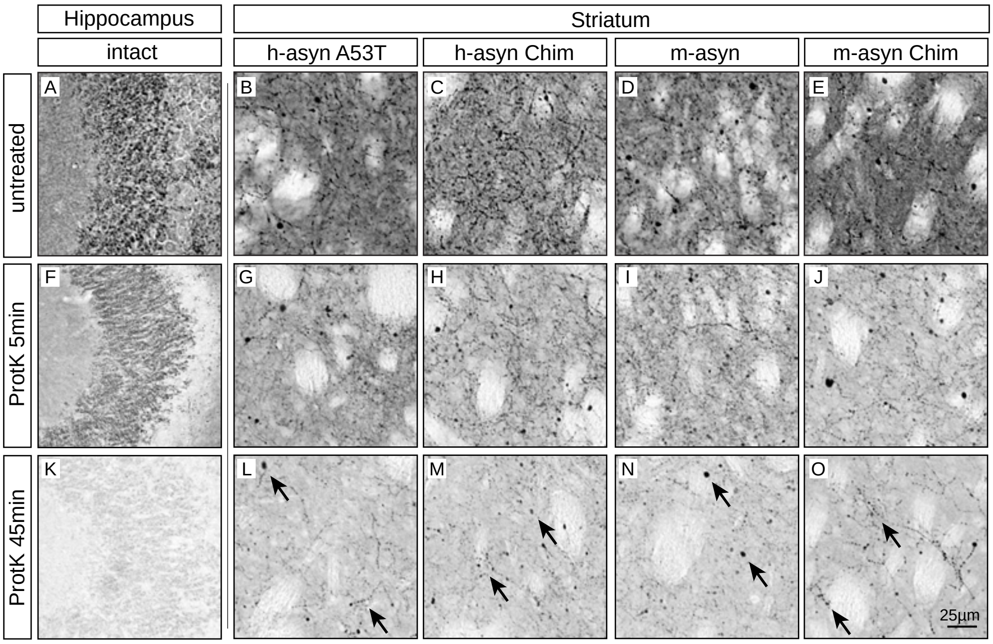
Hippocampus
Striatum
intact
h-asyn A53T
h-asyn Chim
m-asyn
m-asyn Chim
untreated
ProtK 5min
ProtK 45min
A
B
C
D
E
F
G
H
I
J
K
L
M
N
O
25µm

Figure S3

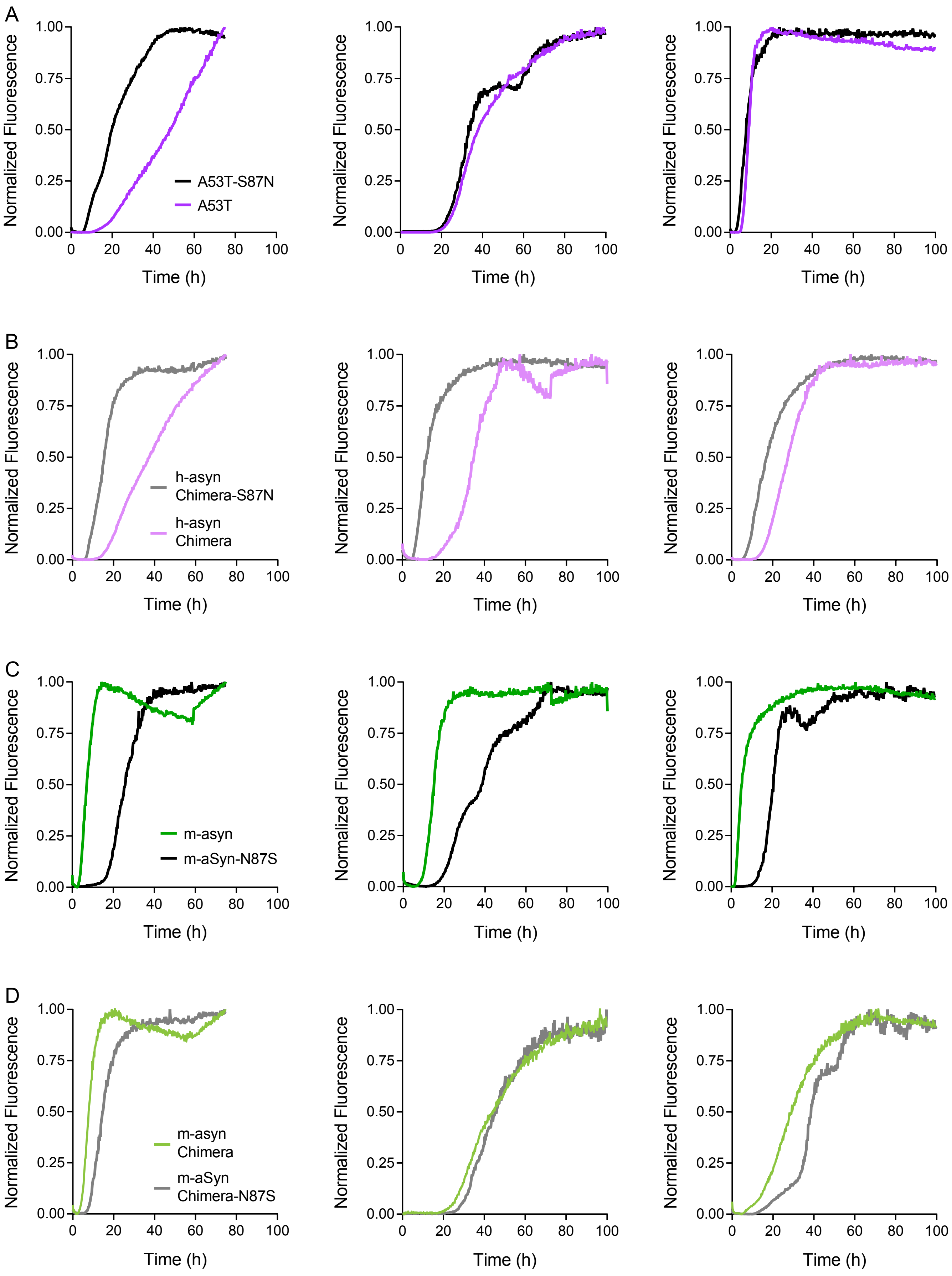
A
B
C
D
Normalized Fluorescence
Time (h)
A53T-S87N
A53T
h-asyn Chimera-S87N
h-asyn Chimera
m-asyn
m-aSyn-N87S
m-asyn Chimera
m-aSyn Chimera-N87S
